# Harnessing the Evolution of Proteostasis Networks to Reverse Cognitive Dysfunction

**DOI:** 10.1101/2025.02.28.640897

**Authors:** Lucas C. Reineke, Ping Jun Zhu, Udit Dalwadi, Sean W. Dooling, Yuwei Liu, I-Ching Wang, Sara Young-Baird, James Okoh, Santosh Kumar Kuncha, Hongyi Zhou, Akshara Kannan, Hyekyung Park, Nicolas A. Debeaubien, Tristan Croll, D. John Lee, Christopher Arthur, Thomas E. Dever, Peter Walter, Jin Chen, Adam Frost, Mauro Costa-Mattioli

## Abstract

The integrated stress response (ISR) is a highly conserved network essential for maintaining cellular homeostasis and cognitive function. Here, we investigated how persistent ISR activation impacts cognitive performance, primarily focusing on a PPP1R15B^R658C^ genetic variant associated with intellectual disability. By generating a novel mouse model that mimics this human condition, we revealed that this variant destabilizes the PPP1R15B•PP1 phosphatase complex, resulting in chronic ISR activation, impaired protein synthesis, and deficits in long-term memory. Importantly, we found that the cognitive and synaptic deficits in *Ppp1r15b*^R658C^ mice are directly due to ISR activation. Leveraging insights from evolutionary biology, we characterized DP71L, a viral orthologue of PPP1R15B, through detailed molecular and structural analyses, uncovering its mechanism of action as a potent pan-ISR inhibitor. Remarkably, we found that DP71L not only buffers cognitive decline associated with a wide array of conditions—including Down syndrome, Alzheimer’s disease and aging—but also enhances long-term synaptic plasticity and memory in healthy mice. These findings highlight the promise of utilizing evolutionary insight to inform innovative therapeutic strategies.

## Introduction

A promising frontier for medical innovation lies in the integrated stress response (ISR), a highly conserved signaling pathway that plays a crucial role in maintaining cell health by reprogramming protein synthesis^1,2^. The ISR originally emerged as a basic mechanism for simple organisms to sense nutrient availability. Over time, it has evolved into a more sophisticated and complex network that helps higher organisms cope with various environmental stresses. In yeast, the ISR is primarily controlled by the kinase Gcn2 and the protein phosphatase 1/Glc7 (PP1/Glc7)^3,4^. In contrast, vertebrates have four distinct ISR kinases—GCN2, PERK, PKR, and HRI—each designed to respond to specific stress signals. Moreover, mammals possess two phosphatase complexes, each containing PP1 bound to either PPP1R15A (GADD34) or PPP1R15B (CReP), regulatory subunits that render the PP1-catalytic subunit specific to the substrate^5,6^. The evolutionary diversification seen in higher organisms has equipped them to effectively adapt and respond to a wide range of environmental and physiological challenges.

Co-evolution between host defense mechanisms and viral strategies has optimized the ISR in mammals to respond to stress. For instance, PKR, the most recently evolved member of the ISR kinase family, plays a critical role in halting protein synthesis during viral infections, protecting cells from viruses that rely on double-stranded RNA for replication^7^. As a countermeasure, viruses have developed sophisticated proteins that specifically counteract the ISR^8^, thereby facilitating viral protein synthesis. This dynamic co-evolution underscores the ongoing arms race between host defense mechanisms and viral strategies for replication and evasion. The potential of these proteins may extend beyond their role in viral propagation as they can be engineered and repurposed to promote health, potentially leading to innovative treatments for ISR-related pathologies.

At the heart of the ISR is the eIF2•eIF2B regulatory node, which modulates the formation of the eIF2•GTP•methionyl-initiator tRNA ternary complex (TC), a key step for initiating protein synthesis. The ISR kinases phosphorylate eIF2 (eIF2-P), which inhibits the activity of eIF2’s guanine nucleotide exchange factor, eIF2B. As a result, eIF2B’s ability to facilitate the exchange of GDP for GTP on eIF2 is reduced, leading to a reduction in TC formation and a general downregulation of protein synthesis rates^1^. Concurrently, ISR activation enhances the translation of certain mRNAs with upstream open reading frames (uORFs) in their 5’ untranslated regions (5’ UTRs), such as ATF4^9,10^. By downregulating global translation and promoting the synthesis of stress-response proteins, the ISR initiates a transcriptional program aimed at restoring cellular homeostasis. The dephosphorylation of eIF2-P is tightly regulated by two regulatory subunits, PPP1R15A and PPP1R15B, which direct the activity of the PP1 catalytic subunit towards eIF2-P^1^. While PPP1R15B is constitutively expressed, PPP1R15A is induced in response to ISR activation^5,6^. Although most of the research on the ISR has been devoted to the study of the ISR kinases and the regulation of eIF2•eIF2B, the role and precise mechanism of action of the ISR phosphatases is relatively less explored, especially in the context of disease.

In the brain, the ISR’s function evolved beyond stress maintenance, playing a crucial role in regulating long-lasting synaptic plasticity and memory formation. Genetic and pharmacological studies across several animal models, including birds and rodents, have demonstrated that inhibition of the ISR enhances long-term memory, whereas activation of the ISR impairs it^11–21^. More importantly, emerging evidence suggests that persistent ISR activation underlies the long-term memory deficits across a wide array of disorders^1^. However, little is known about the underlying mechanisms by which persistent ISR activation leads to cognitive dysfunction.

To explore this issue, we turned to human genetics, as specific gene variants that activate the ISR have been identified in individuals with intellectual disability^22–27^. Most traditional ISR studies utilize acute, nonphysiological conditions to activate the ISR, such as treatment with tunicamycin, a natural product that inhibits the initial step of glycoprotein synthesis and strongly activates the PERK-branch of the ISR. It remains uncertain whether brain cells *in vivo* encounter such severe insults, or if lower levels of ISR activation are more common. Moreover, we do not yet know if the principles underlying acute ISR activation also apply to chronic ISR activation, highlighting the need for models that mimic the physiologically relevant, persistent ISR activation seen in pathological conditions^1^. Here we i) generated a new model of persistent ISR activation driven by a human genetic variant^22,23^ in the phosphatase cofactor PPP1R15B that closely mimics the human condition of intellectual disability, ii) provided evidence that persistent ISR activation is causally linked to the cognitive dysfunction observed in this model, iii) defined the translational program controlled by persistent ISR activation in the brain, and iv) developed and characterized a novel therapeutic approach based on evolutionary biology that reverses the cognitive and synaptic deficits associated with Down Syndrome, Alzheimer’s disease and aging, all conditions in which ISR activation is a prominent feature. In addition, the accompanying manuscript (Törkenczy *et al.*) leverages this mouse model of persistent ISR activation alongside with multi-omic approaches to create a robust platform for dissecting the molecular mechanisms underlying ISR-mediated cognitive decline, revealing that ISR activation impacts distinct brain cell types differently, identifies critical downstream ISR effectors unique to specific cells, and defines an ISR signature through single-cell analyses that is conserved across models of cognitive dysfunction.

## Results

### The ISR is persistently activated in the brains of *Ppp1r15b*^R658C^ mice

The identification of gene variants that activate the ISR in humans with intellectual disability underscores the importance of the ISR in cognitive processing^22–27^. Some of these variants map to the PP1 binding site of PPP1R15B and are predicted to destabilize the PPP1R15B•PP1 phosphatase complex thereby leading to increased eIF2-P and persistent ISR activation^22,23^. However, direct evidence establishing a causal link between these variants and cognitive dysfunction is missing. We employed CRISPR-mediated gene editing to introduce the human PPP1R15B^R658C^ variant into the mouse *Ppp1r15b* gene, generating *Ppp1r15b*^R658C^ mice (**Figure 1A** and see methods). Like human individuals carrying loss-of-function variants in PPP1R15B^22–24^, *Ppp1r15b*^R658C^ mice exhibit smaller brains (**Figure 1B**). PPP1R15B levels in the brains of *Ppp1r15b*^R658C^ mice were comparable to those found in wild-type (WT) mice (**Figure 1C**). Co-immunoprecipitation experiments revealed that PPP1R15B^R658C^ interacted less effectively with PP1 compared to the WT counterpart (**Figure 1D**). Although WT and mutant PPP1R15B exhibit similar binding to eIF2 (**Figure 1D**), the reduced interaction between PPP1R15B^R658C^ and PP1 impaired the phosphatase complex’s ability to dephosphorylate eIF2, resulting in increased eIF2-P levels in the brain of *Ppp1r15b*^R658C^ mice (**Figure 1E**). Hence, the ISR is persistently activated in the brain of *Ppp1r15b*^R658C^ mice.

**Figure 1.**
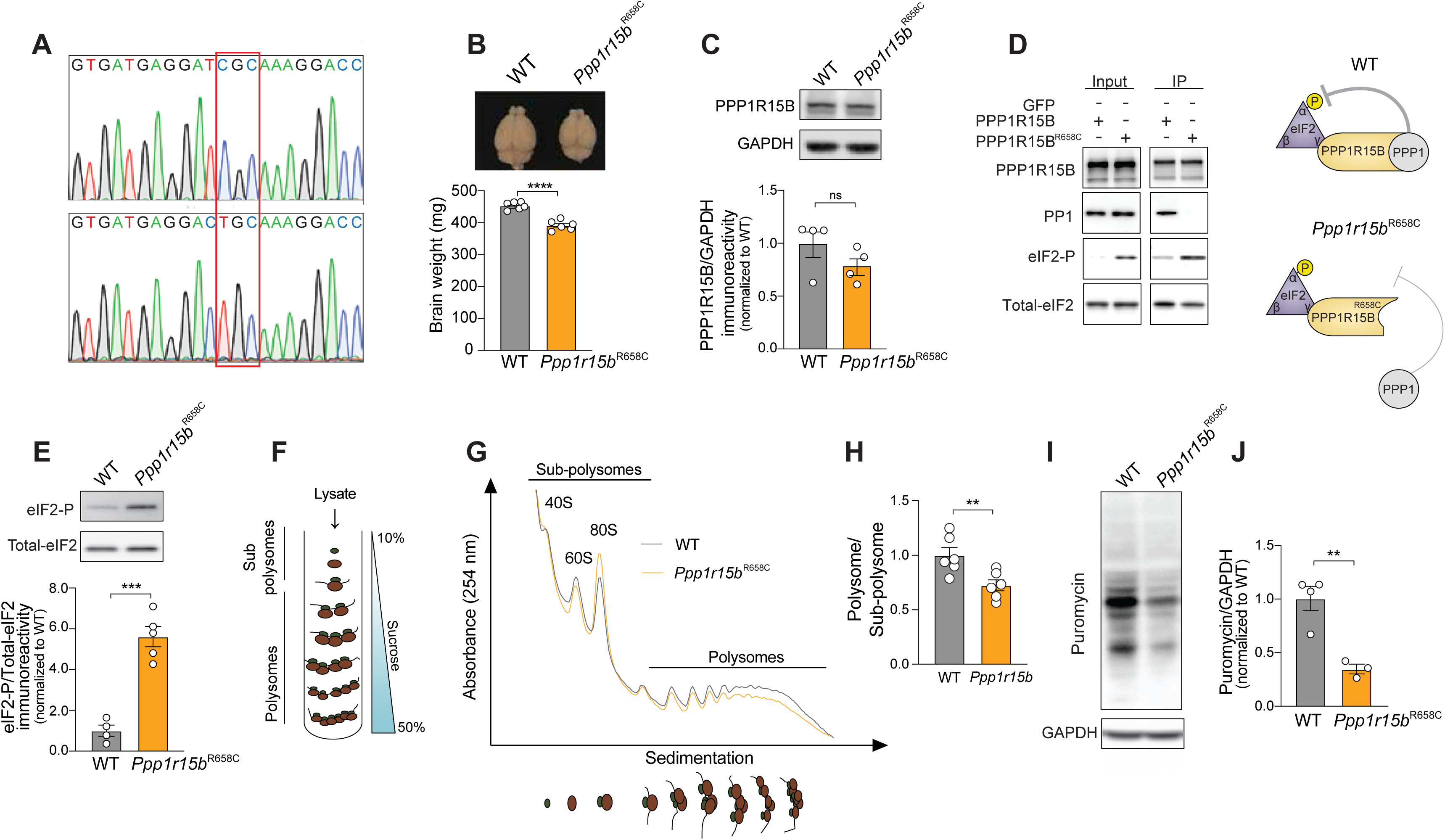
The *Ppp1r15b*^R658C^ variant leads to persistent ISR activation and disrupt protein synthesis in the brain of *Ppp1r15b*^R658C^ mice. (A) Sanger sequencing confirms the generation of *Ppp1r15b*^R658C^ mice carrying the *Ppp1r15b*^R658C^ variant. (B) Comparison of brain size between WT (*n* = 6) and *Ppp1r15b*^R658C^ (*n* = 6) mice (*t* = 7.82, *P* < 0.0001; two-tailed Student’s *t*-test). (C) Representative Western blot (top) and quantification (bottom) of PPP1R15B protein levels in hippocampal extracts from WT (*n* = 4) and *Ppp1r15b*^R658C^ mice (*n* = 4) [*t* = 1.42, *P* = 0.21]. (D) Immunoprecipitation of WT PPP1R15B and mutant PPP1R15B^R658C^ using an GFP antibody followed by Western blots of endogenous PP1 and eIF2 (left). A schematic of the WT and mutant phosphatase complex (right). (E) Representative Western blot (top) and quantification (bottom) of eIF2-P levels in hippocampal extracts from WT (*n* = 4) and *Ppp1r15b*^R658C^ mice (*n* = 5) [*t* = 3.82*, P* < 0.001]. (F) Schematic of polysome profiling sedimentation. After ultracentrifugation, sub-polysomes (40S, 60S, and 80S) and polysomes are separated based on their size. (**G-H**) Averaged polysome profile traces (**G**) and quantification (**H**) of polysome/sub-polysome ratio in the brain of WT and *Ppp1r15b*^R658C^ mice (*n* = 6 per group, *t* = 3.19*, P* < 0.01). (**I-J**) Detection of puromycin incorporation into nascent peptides using an anti-puromycin antibody. A representative Western blot (**I**) and quantification (**J**) in hippocampal extract from WT (*n* = 4) and *Ppp1r15b*^R658C^ mice (*n* = 3) [*t* = 4.73, *P* < 0.01]. Data are mean ± SEM. \**P* < 0.05, \*\**P* < 0.01, \*\*\**P* < 0.001, \*\*\*\**P* < 0.0001).

To assess the impact of sustained ISR activation on protein synthesis, we examined ternary complex levels in the brain of WT and *Ppp1r15b*^R658C^ mice. This complex, which consists of eIF2, GTP, and Met-tRNAi^Met^, is crucial for initiating translation, and is reduced by ISR activation^1^ (**Figure S1A**). When we immunoprecipitated eIF2β, isolated RNA, and performed RT-PCR with primers specific for Met-tRNAi^Met^, we found a reduction in Met-tRNAi^Met^ levels in *Ppp1r15b*^R658C^ mice, indicating that ternary complex levels are reduced in the brains of mutant mice (**Figure S1B-C**). We next examined overall protein synthesis rates by polysome sedimentation in sucrose gradients. This method differentiates mRNAs based on the degree of ribosome association: mRNAs with fewer ribosomes (monosomes) sediment near the top of the gradient, while those with multiple ribosomes (polysomes) sediment toward the bottom (**Figure 1F**). We found decreased polysome and increased monosome (80S) ratios in the brain of *Ppp1r15b*^R658C^ mice (**Figure 1G-H**), characteristic of inhibited protein synthesis at the initiation step^28^. In addition, *in vivo* protein synthesis, measured by puromycin incorporation into nascent polypeptide chains, was also reduced in the brain of *Ppp1r15b*^R658C^ mice (**Figure 1I-J**). In summary, the *Ppp1r15b*^R658C^ variant destabilizes the PPP1R15B•PP1 phosphatase complex, leading to sustained ISR activation and diminished protein synthesis rates in the brain of *Ppp1r15b*^R658C^ mice.

### *Ppp1r15b*^R658C^ mice exhibit deficits in long-term memory and synaptic function due to ISR activation

While individuals with the PPP1R15B^R658C^ variant show intellectual disability^22–24^, direct causal links between this variant and the molecular, cellular, and behavioral changes associated with cognitive dysfunction have yet to be established. Thus, we assessed cognitive function in *Ppp1r15b*^R658C^ mice and WT littermates by examining their long-term contextual fear memory. In this task, mice are exposed to a context (conditioned stimulus; CS) paired with a foot shock (the unconditioned stimulus; US). Twenty-four hours after training, mice were exposed to the CS and fear responses (“freezing behavior”) were measured as an index of the strength of their long-term memory (**Figure 2A**). We found that freezing behavior prior to training was similar in naïve *Ppp1r15b*^R658C^ mice and control littermates. However, *Ppp1r15b*^R658C^ mice exhibited a significant reduction in freezing 24 hr after training, indicating that their long-term memory is impaired (**Figure 2B**). Likewise, late long-term potentiation (LTP), a cellular model of long-term memory^29^, was also impaired in slices from *Ppp1r15b*^R658C^ mice (**Figure 2C**). The impaired late-LTP was not due to deficits in basal synaptic transmission, as input-output plots of field excitatory potentials as a function of presynaptic fiber volley and paired pulse facilitation were comparable between WT and *Ppp1r15b*^R658C^ mice (**Figure S2**). Hence, persistent ISR activation impairs both long-term synaptic and memory processes.

**Figure 2.**
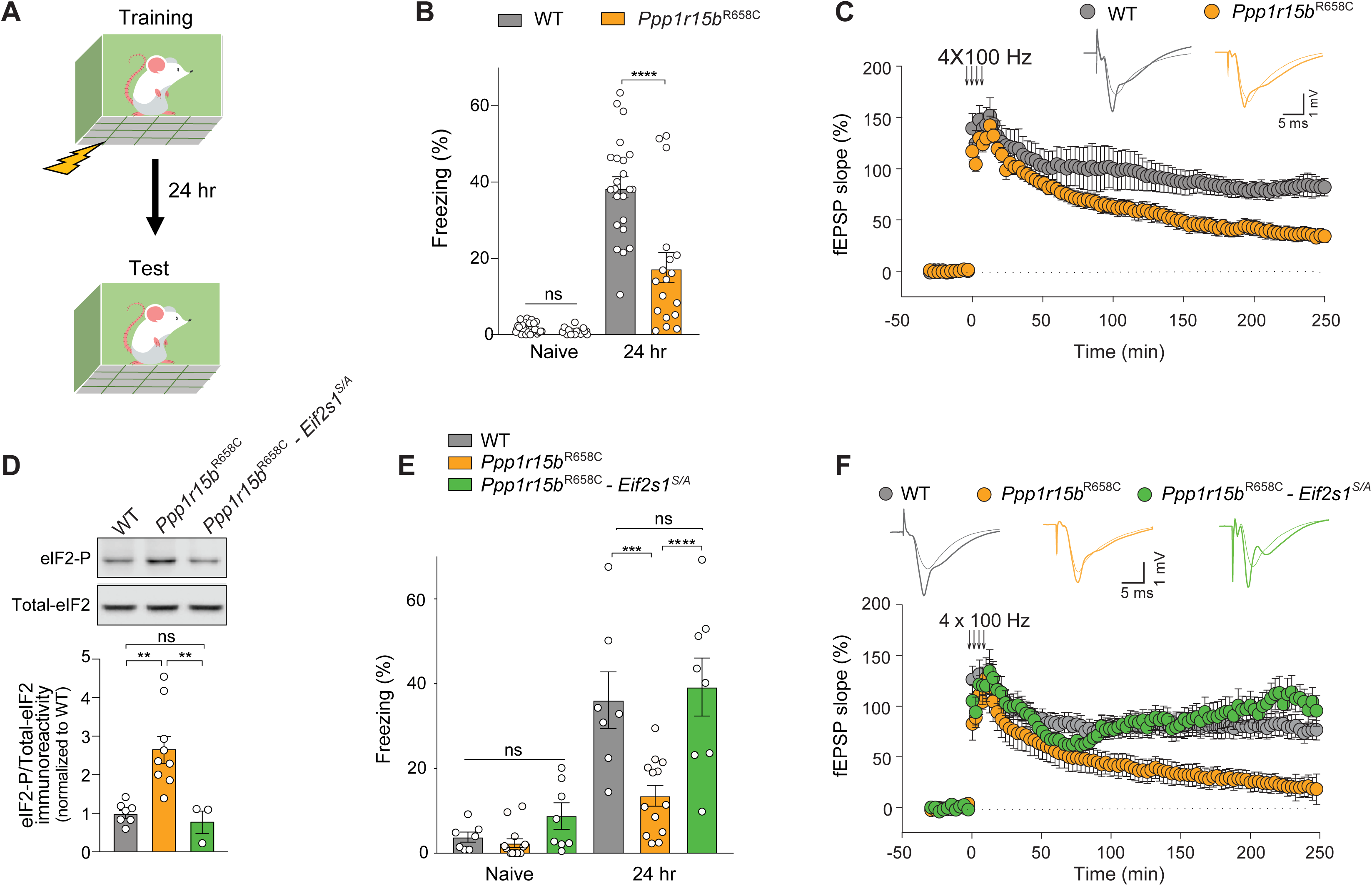
Persistent ISR activation cause deficits in long-term synaptic plasticity and memory in *Ppp1r15b*^R658C^ mice. (A) Schematic of the contextual fear conditioning paradigm. (B) Long-term contextual fear memory in WT (*n* = 23) and *Ppp1r15b*^R658C^ (*n* = 18) mice (*F*3,52 = 60.76, *t* = 5.73, *P* < 0.0001). (C) L-LTP induced by four trains of high frequency stimulation (HFS, 4 x 100 Hz) in WT (*n* = 8) and *Ppp1r15b*^R658C^ (*n* = 12) mice (at 250 min: *t* = 4.14, *P* < 0.001). (D) Representative Western blot (top) and quantification (bottom) of eIF2-P levels in hippocampal extracts from WT (*n* = 7), *Ppp1r15b*^R658C^ mice (*n* = 9) and *Ppp1r15b*^R658C^*-Eif2s1^S/A^* (*n* = 3) [*F*2,16 = 1.86, WT vs. *Ppp1r15b*^R658C^: *t* = 4.25, *P* < 0.01; *Ppp1r15b*^R658C^ vs. *Ppp1r15b*^R658C^*-Eif2s1^S/A^*: *t* = 3.60, *P* < 0.01; WT vs. *Ppp1r15b*^R658C^*- Eif2s1^S/A^*: *t* = 0.39, *P* > 0.99). (E) Long-term contextual fear memory in WT (*n* = 7), *Ppp1r15b*^R658C^ (*n* = 13) and *Ppp1r15b*^R658C^*- Eif2s1^S/A^* mice (*n* = 8) [*F*2,50 = 5.58; WT vs. *Ppp1r15b*^R658C^: *t =* 4.31*, P <* 0.001*; Ppp1r15b^R658C^* vs. *Ppp1r15b*^R658C^*- Eif2s1^S/A^*: *t =* 5.12*, P* < 0.0001; WT vs. *Ppp1r15b*^R658C^*- Eif2s1^S/A^*: *t =* 0.54*, P >* 0.99). (F) L-LTP induced by four trains of high frequency stimulation (HFS, 4 x 100 Hz) in WT (*n* = 9) and *Ppp1r15b*^R658C^ (*n* = 5) mice and *Ppp1r15b*^R658C^*- Eif2s1^S/A^*(*n* = 8) [at 250 min: *F*2,19 = 8.69; WT vs. *Ppp1r15b*^R658C^: *t =* 3.18*, P* < 0.01; *Ppp1r15b*^R658C^ vs. *Ppp1r15b*^R658C^*- Eif2s1^S/A^*: *t* = 4.09, *P* < 0.001; WT vs. *Ppp1r15b*^R658C^*- Eif2s1^S/A^*: *t* = 1.15, *P* = 0.264). Data are mean ± SEM. \**P* < 0.05, \*\**P* < 0.01, \*\*\**P* < 0.001, \*\*\*\**P* < 0.0001).

Given that a delicate interplay between excitatory and inhibitory synaptic activity is essential for long-term memory formation^30^, we next examined if this balance was altered in the brain of *Ppp1r15b*^R658C^ mice. To this end, we measured inhibitory and excitatory synaptic transmission in the hippocampus of WT and *Ppp1r15b*^R658C^ mice. Whole-cell recordings revealed that both the frequency and amplitude of miniature excitatory postsynaptic currents (mEPSCs) were comparable between *Ppp1r15b*^R658C^ and WT mice (**Figure S3A-C**). In contrast, we found that the frequency (but not the amplitude) of miniature inhibitory postsynaptic currents (mIPSCs) was increased in *Ppp1r15b*^R658C^ mice (**Figure S3D-F**). These findings indicate that persistent ISR activation selectively enhances inhibitory synaptic transmission in *Ppp1r15b*^R658C^ mice.

Finally, to determine whether the deficits in long-term memory and synaptic function in *Ppp1r15b*^R658C^ mice are due to ISR activation, we crossed *Ppp1r15b*^R658C^ mice with mice in which one of the alleles of the *Eif2s1* gene (eIF2) is mutated to alanine (*Eif2s1^S/A^*) and cannot be phosphorylated. This cross led to normalized levels of eIF2-P (**Figure 2D**). Remarkably, resetting the ISR fully rescued the cognitive deficits (**Figure 2E**), restored LTP (**Figure 2F**) and corrected the increased inhibitory synaptic transmission in *Ppp1r15b*^R658C^ mice (**Figure S3G-H**). Our results demonstrate that the long-term memory and synaptic plasticity deficits caused in *Ppp1r15b*^R658C^ mice result from persistent ISR activation.

### Persistent ISR activation reprograms the translational landscape in the brain of *Ppp1r15b*^R658C^ mice

The ISR plays a crucial role in maintaining cell health by regulating protein synthesis. To determine the nature of the proteins whose synthesis is altered in the brain due to persistent ISR activation, we performed ribosome profiling (Ribo-Seq), a powerful tool for monitoring mRNA translation *in vivo* ^31^. The *Ppp1r15b*^R658C^ model offers a unique opportunity to identify selective ISR-mediated gene expression changes in the brain, including at the single-cell level (see accompanying manuscript, Törkenczy *et al.*), resulting solely from sustained ISR activation without the confounding effects of other dysregulated pathways often present in mouse models with ISR activation (e.g., Alzheimer’s disease, traumatic brain injury, aging, and Down syndrome)^32–34^. In addition, unlike studies that typically use acute ISR stressors, here we mapped the genome-wide translational landscape resulting from persistent ISR activation in the brain. Briefly, we sequenced ribosome-protected mRNA fragments in WT and *Ppp1r15b*^R658C^ mice. Concurrently, we isolated total RNA and subjected it to RNA-Seq. We found that the ribosome profiling was highly reproducible between samples (**Figure S4A**). As anticipated, we found numerous genes to be transcriptionally and translationally dysregulated in the brain of *Ppp1r15b*^R658C^ mice (**Figure 3A**). Gene set enrichment analysis revealed an enrichment of genes linked with the unfolded protein response (UPR, which shares its PERK-branch with the ISR^1,35^, **Figure 3B**). Given that we did not observe changes in the activation of the UPR sensors (IRE1, ATF6 and PERK; **Figure S5**), the gene enrichment observed in the brain of *Ppp1r15b*^R658C^ mice likely represents ISR activation. Moreover, we found an enrichment in ATF4 target genes (**Figure 3A, 3C**), which we confirmed by qRT-PCR (**Figure 3D**). Please note that in the accompanying manuscript (Törkenczy *et al.*), we define a comprehensive ISR signature at the single-cell level that extends beyond ATF4 targets.

**Figure 3.**
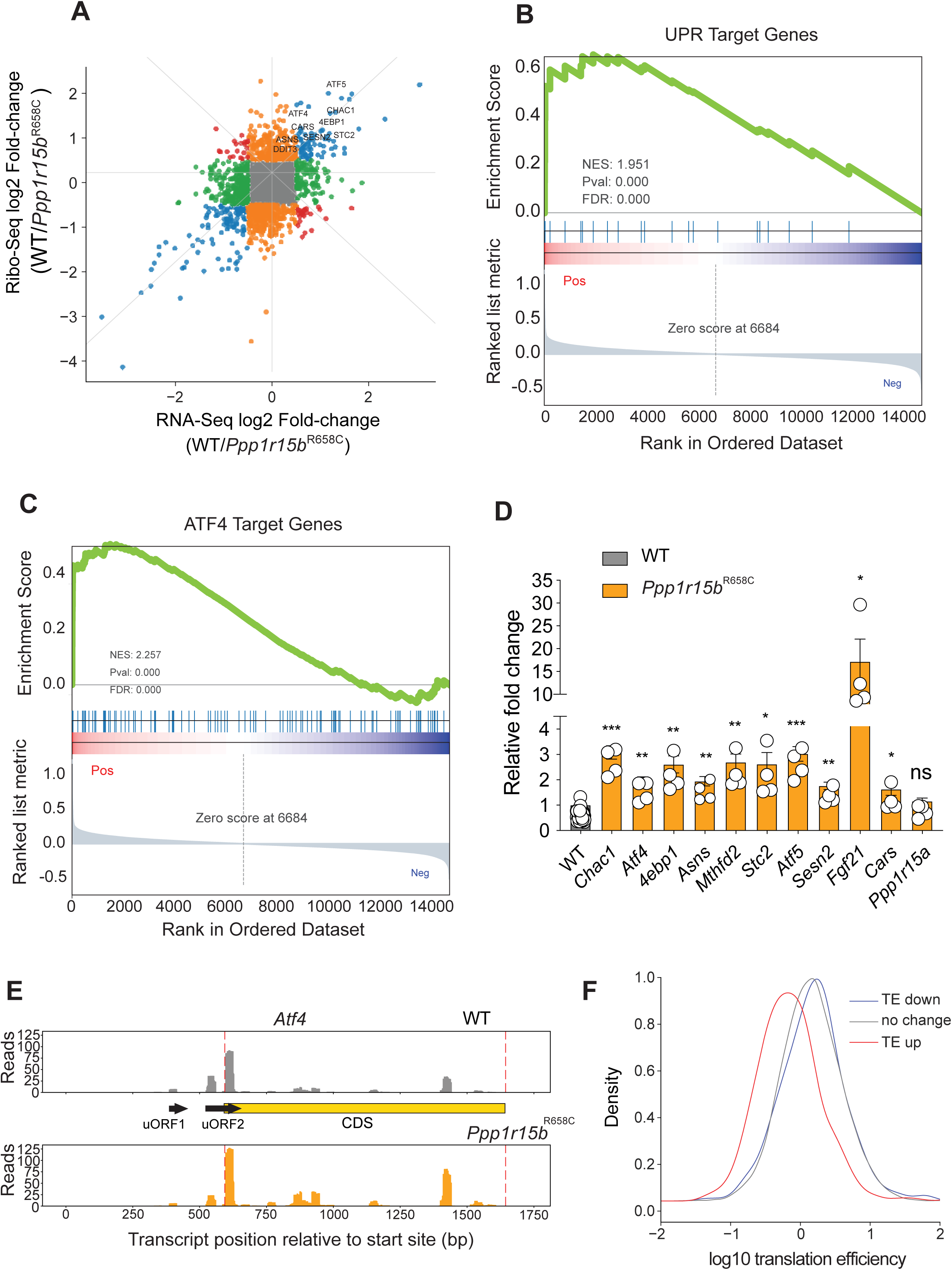
Genome-wide translatome profiling of persistent ISR activation in the brain of *Ppp1r15b*^R658C^ mice. (A) Comparison of translatome and transcriptome analyses derived from ribosome profiling and RNA-seq in the brains of WT (*n* = 3) and *Ppp1r15b*^R658C^ (*n* = 3) mice. (B) Gene set enrichment analysis reveals an increase in unfolded protein response (UPR) genes in the brain of *Ppp1r15b*^R658C^ mice. (C) Gene set enrichment analysis reveals an increase in ATF4 target genes in the brain of *Ppp1r15b*^R658C^ mice. (D) Quantitative PCR for ATF4 target genes in WT (*n* = 4) and *Ppp1r15b*^R658C^ (*n* = 4) mice (*Chac1*: *t* = 7.75, *P* < 0.001; *Atf4*: *t =* 4.41*, P* < 0.01*; Eif4ebp1*: *t =* 4.67*, P* < 0.01*; Asns*: *t =* 4.86*, P* < 0.01*; Mthfd2*: *t =* 5.11*, P* < 0.01*; Stc2 t =* 3.46*, P <* 0.05*; Atf5: t =* 6.66*, P <* 0.001*; Sesn2*: *t =* 4.76*, P* < 0.01*; Fgf21*: *t =* 3.26*, P <* 0.05*; Cars*: *t =* 2.50*, P <* 0.05*; Ppp1r15a*: *t =* 1.13*, P =* 0.30). (E) Ribosome occupancy along the *ATF4* mRNA in WT and *Ppp1r15b*^R658C^ mice. (F) Changes in translation efficiency (TE) in WT and *Ppp1r15b*^R658C^ mice, with mRNAs categorized based on their TE (high, low or unchanged). Data are mean ± SEM. \**P* < 0.05, \*\**P* < 0.01, \*\*\**P* < 0.001, \*\*\*\**P* < 0.0001).

To determine the impact of sustained ISR activation on gene-specific translation, we evaluated translational efficiency (TE) by determining the reads per kilobase of transcript per million mapped reads (RPKM) of coding sequences (CDS) in ribosome profiling versus the RPKM in exons from RNA- seq (RPKMribosome profiling/RPKMRNA-seq) in the brain of WT and *Ppp1r15b*^R658C^ mice. Our analysis revealed that translation efficiency of over 174 mRNAs selectively changed, with 49 increasing and 125 decreasing in translation efficiency in the brain of *Ppp1r15b*^R658C^ mice (**Figure 3A, Figure S4B-C, Table I**). Although a small subset of mRNAs whose translation was increased upon ISR activation have been reported to contain uORFs in their 5’UTR^36^, we found that the length of the 5’ UTR, CDS, or 3’ UTR did not signifcantly influence changes in translation efficiency resulting from sustained ISR activation (**Figure S6A-C**). In addition, we found that translation of mRNAs carrying a 5’TOP-like motif, a cis-regulatory regulatory element in the 5’UTR of some mRNAs that regulate translation rates^37,38^, was not up-regulated in the brain of *Ppp1r15b*^R658C^ mice (**Figure S6D**). Moreover, the translation of mRNAs associated with proliferation (E2F1-driven) and lipid metabolism (SREBF1-driven), previously reported to be controlled by the ISR^39^, show no differences between WT and *Ppp1r15b*^R658C^ mice (**Figure S7A-B**).

In contrast, we found a significant increase in ribosomal occupancy at the ATF4 mRNA’s canonical AUG start site and along the ATF4 CDS, coupled with a decreased ribosome presence at the inhibitory 5’UTR ORF2, indicative of enhanced translation of ATF4 in *Ppp1r15b*^R658C^ mice (**Figure 3E** and **Figure S7C**). Interestingly, while we also found increased ribosomal occupancy on the *Ddit3/*CHOP mRNA, another key ISR target, its downstream target genes did not show corresponding upregulation in the brain of *Ppp1r15b*^R658C^ mice (**Figure S8A-B**). Furthermore, our analysis revealed that mRNAs with higher translation efficiency are more susceptible to the translation inhibition caused by persistent ISR activation, while those with lower TE tend to be resistant to ISR activation (**Figure 3F**). Thus, persistent ISR activation in the brain of *Ppp1r15b*^R658C^ mice does not uniformly reduce the translation of all mRNAs.

Finally, acute ISR activation triggers an increase in PPP1R15A expression at both transcriptional and translational levels^6,40^, serving as a feedback mechanism to mitigate ISR activation. Surprisingly, we found no significant differences in PPP1R15A protein levels, mRNA levels, or ribosome occupancy (**Figure 3D** and **Figure S9A-C**) in the brains of *Ppp1r15b*^R658C^ mice compared to their control counterparts. Together, these findings support the notion that sustained ISR activation reprograms gene expression without triggering the typical (PPP1R15A/GADD34) feedback mechanism associated with an acute ISR response.

### DP71L evolved as a potent repressor of the ISR

Given that disruption of PPP1R15B function causes cognitive dysfunction due to ISR activation (**Figure 1-2**), we hypothesized that PPP1R15B-like molecules with enhanced activity might enhance long-term memory and serve as potential therapies. To explore this possibility, we turned to evolutionary biology, which has already yielded several disruptive innovations. Using comparative genomics, we identified 391 proteins across animals and viruses that shared PP1- and eIF2-binding motifs analogous to those found in PPP1R15B (**Figure 4A** and **Table II**). Among these, DP71L, a protein from African swine fever virus^41,42^, which consists solely of PP1- and eIF2-binding motifs, stood out as the smallest protein in the family. We found that DP71L effectively suppresses the ISR in mammalian cells regardless of the ISR kinase that is activated (see **Figure 4B-F**). Hence, DP71L behaves as a potent pan-ISR suppressor.

**Figure 4.**
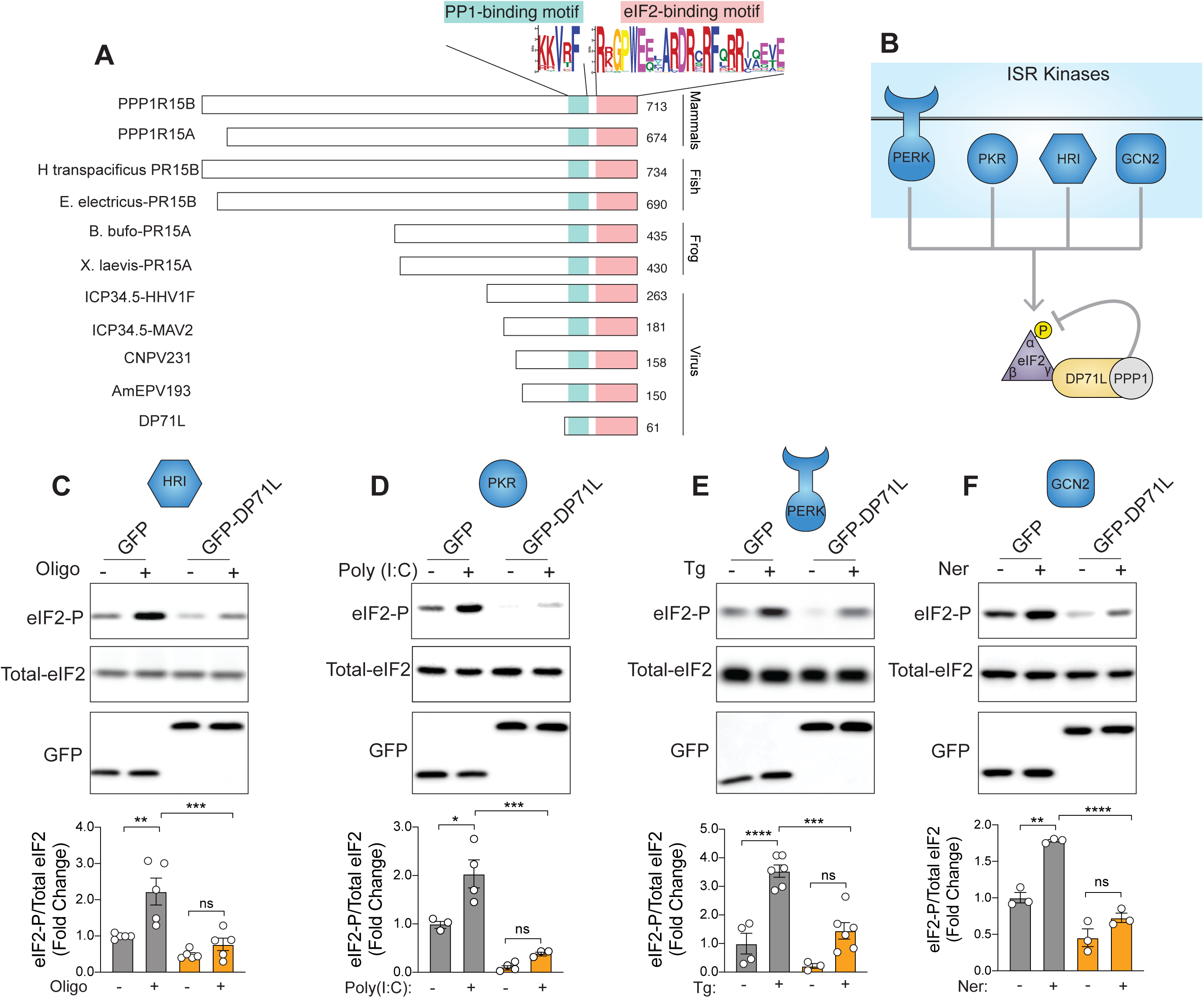
Protein alignment across diverse kingdoms of life reveals DP71L as a potent pan-ISR inhibitor. (A) Schematic of protein alignment showing PP1- and eIF2-binding motifs across various species. Consensus PP1- and eIF2-binding motif sequences are shown. (B) Overview of ISR kinases and the mechanism by which DP71L represses the ISR. (**C-E**) Cells expressing either GFP or GFP-DP71L were treated with the specific chemical inducer for each ISR kinase. Representative Western blot (top) and quantification (bottom) of eIF2-P levels upon activation of HRI (**C**, *F*3,16 *=* 7.14, *n* = 5 per group; GFP+Vehicle vs. GFP+Oligo: *t* = 4.22, *P* < 0.01; GFP-DP71L+Vehicle vs. GFP-DP71L+Oligo: *t* = 0.98, *P* > 0.99; GFP+Oligo vs. GFP-DP71L+Oligo: *t* = 4.17, *P* < 0.01), PKR [**D**, *n* = 3-4 per group, *F*3,10 *=* 9.80; GFP+Vehicle vs. GFP+Poly(I:C): *t* = 4.16, *P* < 0.05; GFP-DP71L+Vehicle vs. GFP-DP71L+Poly(I:C): *t* = 1.15, *P* > 0.99; GFP+Poly(I:C) vs. GFP- DP71L+Poly(I:C): *t* = 6.13, *P* < 0.001], PERK (**E,** *n* = 3-6 per group, *F*3,15 *=* 1.90, GFP+Vehicle vs. GFP+Tg: *t* = 5.52, *P* < 0.0001; GFP-DP71L+Vehicle vs. GFP-DP71L+Tg: *t* = 2.90, *P* = 0.06; GFP+Tg vs. GFP-DP71L+Tg: *t* = 5.99, *P* < 0.001) and GCN2 (**F,** *n* = 3 per group, *F*3,8 *=* 0.67; GFP+Vehicle vs. GFP+Ner: *t* = 6.29, *P* < 0.01; GFP-DP71L+Vehicle vs. GFP-DP71L+Ner: *t* = 2.20, *P* < 0.35; GFP+Ner vs. GFP-DP71L+Ner: *t* = 9.27, *P* < 0.001). Tg = thapsigargin, Oligo = oliogomycin, Ner = neratinib, Poly:IC = polyinosinic-polycytidylic acid: a synthetic double strand RNA. Data are mean ± SEM. \**P* < 0.05, \*\**P* < 0.01, \*\*\**P* < 0.001, \*\*\*\**P* < 0.0001).

In contrast to its larger mammalian counterparts PPP1R15A and PPP1R15B, which have additional N-terminal motifs aiding in eIF2 binding^24,43^, DP71L’s simpler structure prompted us to investigate its unique ISR suppression abilities. Head-to-head comparison between DP71L and a PPP1R15B protein of the same size (ΔPPP1R15B containing only the PP1- and eIF2-binding motifs, **Figure 5A**), revealed that DP71L suppresses the ISR significantly more efficiently than ΔPPP1R15B (**Figure 5B**, compare lane 2 to 4). Moreover, we found that DP71L, despite lower expression levels, pulled down PP1 more efficiently than ΔPPP1R15B (**Figure 5C**), indicating that DP71L binds more strongly to PP1. Note that both DP71L and ΔPPP1R15B pulled down similar amounts eIF2 (**Figure 5C**).

**Figure 5.**
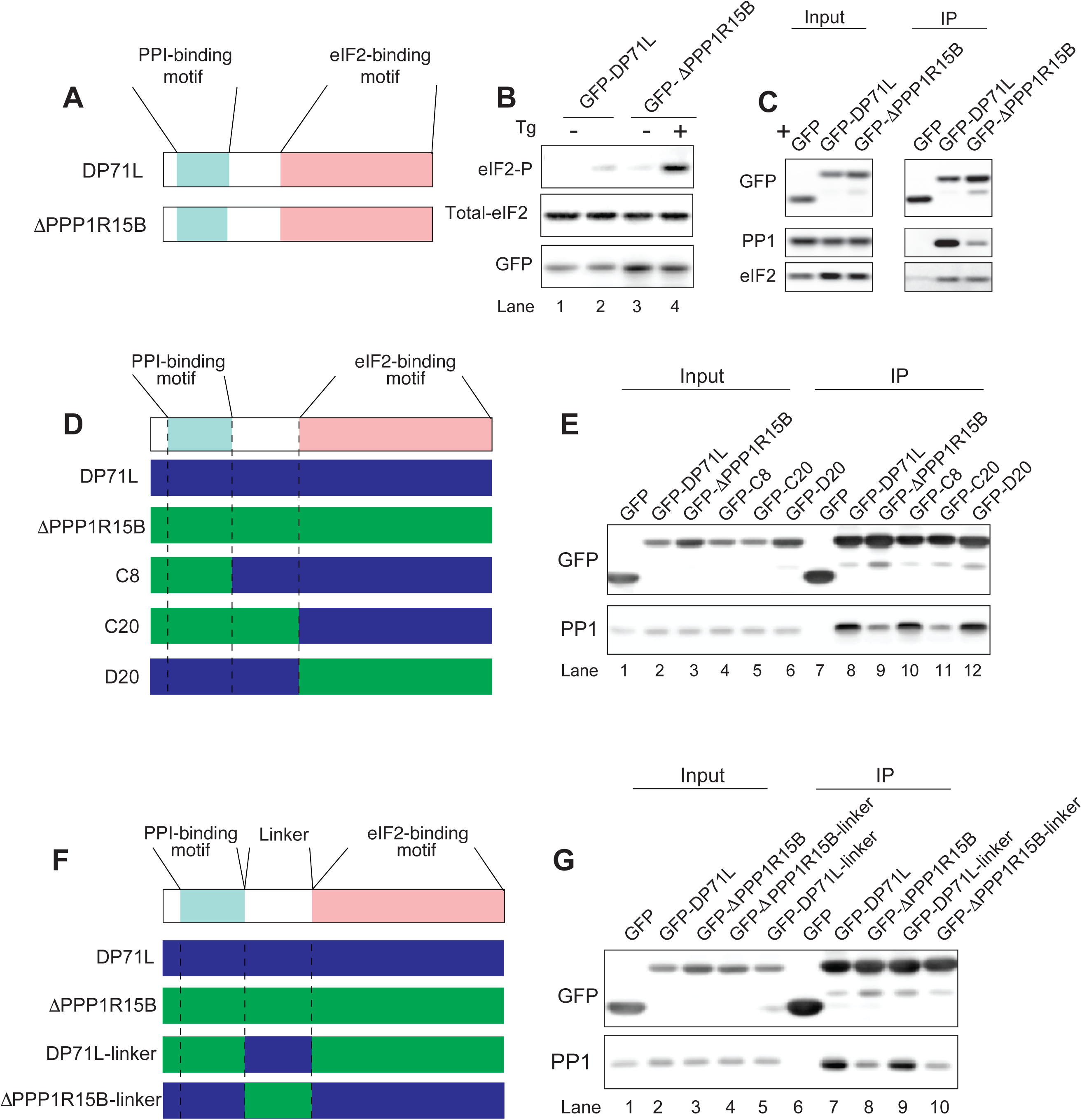
Molecular dissection of DP71L function. (A) Schematic comparison of DP71L and ΔPPP1R15B, highlighting their PP1- and eIF2-binding motifs. (B) Representative Western blot of eIF2-P levels following expression of either GFP-DP71L or GFP- ΔPPP1R15B. (C) Immunoprecipitation of GFP-DP71L or GFP-ΔPPP1R15B using a GFP antibody, followed by Western blot against endogenous PP1 or eIF2. (D) Schematic of the different GFP-DP71L/GFP-ΔPPP1R15B chimeras. (E) Immunoprecipitation of the different chimeras with a GFP antibody, followed by Western blotting for endogenous PP1. (F) Schematic of various GFP-DP71L and GFP-ΔPPP1R15B chimeras. (G) Immunoprecipitation of the different chimeras using a GFP antibody, followed by Western blotting for endogenous PP1.

To identify the domain of DP71L responsible for its enhanced binding to PP1, we generated chimeric proteins by exchanging motifs between DP71L and ΔPPP1R15B (**Figure 5D**). We found that compared to ΔPPP1R15B, ΔPPP1R15B chimeras carrying the DP71L-linker region (the region between the PP1- and eIF2-binding motifs) exhibit better binding to PP1 regardless of whether they also contained the eIF2-binding motif (**Figure 5E**, D20, compare lanes 9 to 12) or the PP1-binding motif of PPP1R15B (**Figure 5E**, C8, compare lanes 9 to 10). Conversely, compared to DP71L, the DP71L chimera containing ΔPPP1R15B-linker region binds less efficiently to PP1 (**Figure 5E**, C20, compare lanes 8 to 11). Moreover, a DP71L chimera containing its linker region binds to PP1 with similar affinity as WT DP71L, irrespective of whether they contain their native PP1-binding motif (**Figure 5E**, C8, compare lanes 8 to 10) or the eIF2-binding motif (**Figure 5E**, D20, compare lanes 8 to 12) of PPP1R15B. Taken together, these results indicate that the sequence linking the PP1 and eIF2 binding motifs (“Linker”) determines the differential binding of these proteins to PP1. To test this directly, we swapped only the linker regions of ΔPPP1R15B and DP71L (**Figure 5F**). We found that the DP71L chimera carrying the ΔPPP1R15B-linker region exhibits reduced binding to PP1 compared to DP71L (**Figure 5G**, compare lanes 7 to 10). The opposite is also true, compared to ΔPPP1R15B, a ΔPPP1R15B chimera carrying the DP71L-linker region binds more efficiently to PP1 (**Figure 5G**, compare lane 8 to 9). Hence, the linker region of DP71L confers its enhanced PP1-binding ability, indicating that extended residues, beyond the previously described four amino acids defining the well-characterized RVxF motif^42,44–46^, participate in effective PP1 binding.

To further elucidate the mechanism by which DP71L suppresses the ISR, we solved the structure of the phosphatase-bound complex by cryo-electron microscopy (cryo-EM; **Figure S10**). Based on earlier studies of the mammalian phosphatase complexes^47^, we stabilized the PP1•DP71L•eIF2 complex with G-actin and DNase I. In brief, we obtained a ∼ 3.1 Å resolution map of the complex, including detailed information about the linker region of DP71L (**Figure 6A**). By modeling the ΔPPP1R15B sequence into the density map of the DP71L complex, we compared the linker regions of DP71L and ΔPPP1R15B. Consistent with the higher binding affinity of DP71L to PP1, we found the linker of DP71L exhibited novel interactions with PP1. Notably, Asp27 in the DP71L linker forms a salt bridge with Arg74 of PP1, an interaction that is absent between ΔPPP1R15B and PP1. In addition, the DP71L linker lacked a bulge created by the presence of Gly654 in ΔPPP1R15B, which decreases the local interaction surface area from 1562 Å for DP71L-PP1 to 1410 Å for PPP1R15B-PP1 (**Figure 6B**), as determined by the PDBePISA tool^48^. To determine if these modifications in the linker of DP71L indeed enhance its binding to PP1, we introduced selective mutations in the linker of DP71L to mimic ΔPPP1R15B: briefly, we replaced Asp27 with a Ser (as in ΔPPP1R15B) and added the equivalent of Gly654, which increased the bulge size (DP71L-4X). OpenFold modeling^49^ predicted that DP71L-4X would bulge out and disrupt the salt bridge, assuming a structure similar to ΔPPP1R15B (**Figure 6C**). Indeed, pulldown experiments showed DP71L-4X exhibited weaker binding to PP1 compared to DP71L (**Figure 6D**), confirming the importance of these residues in the binding between DP71L and PP1. Conversely, we engineered gain-of-function mutations in ΔPPP1R15B to emulate DP71L by introducing the salt bridge and removing Gly654, which reduces the size of the bulge (ΔPPP1R15B-3X). OpenFold predicted that the ΔPPP1R15B-3X mutation would improve binding to PP1 by increasing the buried surface area and re-introducing the salt bridge (**Figure 6E**), a prediction that was confirmed by our pull-down experiments (**Figure 6F**). Together, the cryo-EM and biochemical studies explain how DP71L potently suppresses the ISR, relying on a crucial salt bridge and the absence of a bulge as compared to ΔPPP1R15B to enhance PP1 binding.

**Figure 6.**
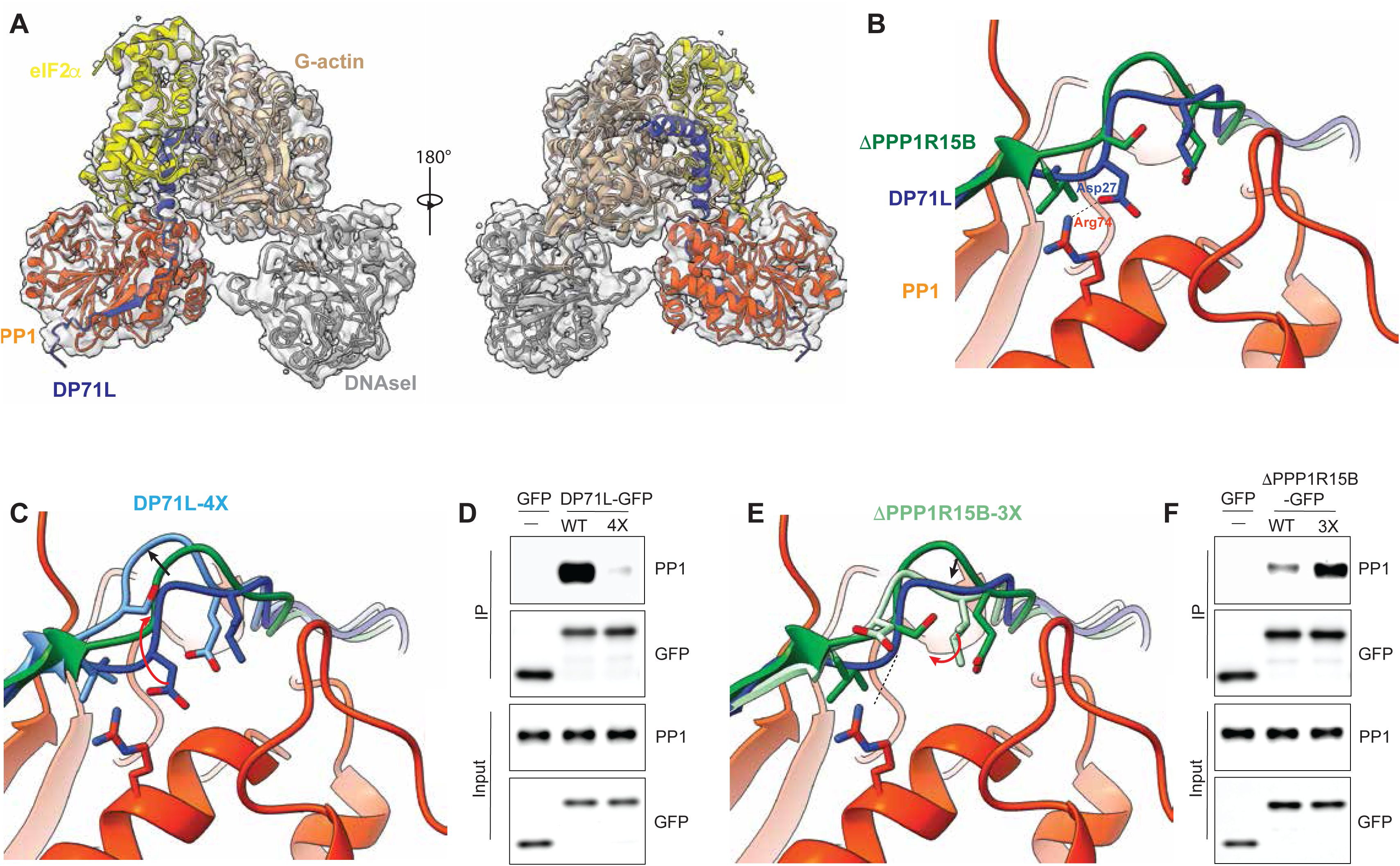
Structural characterization of DP71L function. (A) A 3.1 Å electron density map of the PP1•DP71L•eIF2α•G-actin complex (left), and the same map rotated 90°. DP71L is colored blue, PP1 is orange, eIF2α is yellow, and G-actin is copper. (B) Zoomed view of the linker region of DP71L and PPP1R15B, which was modeled into the cryo-EM map (green). (C) Structural modeling a DP71L mutant (4X) predicts an increase the size of the bulge and disruption of the hydrogen bonding, likely leading to decreased binding affinity for PP1. (D) Immunoprecipitation of WT GFP-DP71L and 4X GFP-DP71L with an GFP antibody, followed by Western blotting for endogenous PP1. (E) Structural modeling a mutant ΔPPP1R15B (3X) predicts a decrease in the size of the bulge and the formation of a new hydrogen bond, potentially enhancing binding to PP1. (F) Immunoprecipitation of WT ΔPPP1R15B and 3X ΔPPP1R15B with an GFP antibody, followed by Western blotting for endogenous PP1.

### DP71L reverses the long-term memory deficits in several models of cognitive disorders by enhancing synaptic plasticity

Adeno-associated virus (AAV)-based gene therapy is one of the most well-characterized and widely used vectors in gene therapy^50^. The size of the therapeutic gene is crucial in AAV-based gene therapy because the vector has a limited packaging capacity, which includes the therapeutic gene and essential regulatory elements. Thus, using compact gene effectors is essential for maximizing therapeutic potential of AAVs. Given that i) DP71L is a potent pan-ISR inhibitor (**Figure 4C-F**), ii) DP71L is notably small, being the smallest among phosphatase cofactors (**Figure 4A** and **Table II**), and iii) the ISR is activated in the brain of a broad spectrum of cognitive disorders^1^, we wondered whether AAV-mediated DP71L expression could be sufficient to reverse long-term memory deficits across different disease classes.

We first investigated this possibility in the context of Down syndrome, the most common genetic cause of intellectual disability^51^, which is characterized by ISR activation^52^. As previously reported, we found that Ts65Dn mice, a well-established model of Down syndrome that recapitulates the cognitive deficits of the human syndrome, showed impaired long-term fear memory (**Figure 7A**). Remarkably, a single injection of GFP-DP71L into the brain of Ts65Dn mice suppressed the ISR, as determined by reduced eIF2-P levels (**Figure S11A**) and significantly reversed their long-term memory deficits (**Figure 7A**). Furthermore, DP71L-treatment restored the deficits in long-lasting synaptic plasticity in the brains of Ts65Dn mice (**Figure 7B**). Thus, DP71L-mediated suppression of the ISR reverses the cognitive and synaptic deficits associated with Down syndrome.

**Figure 7.**
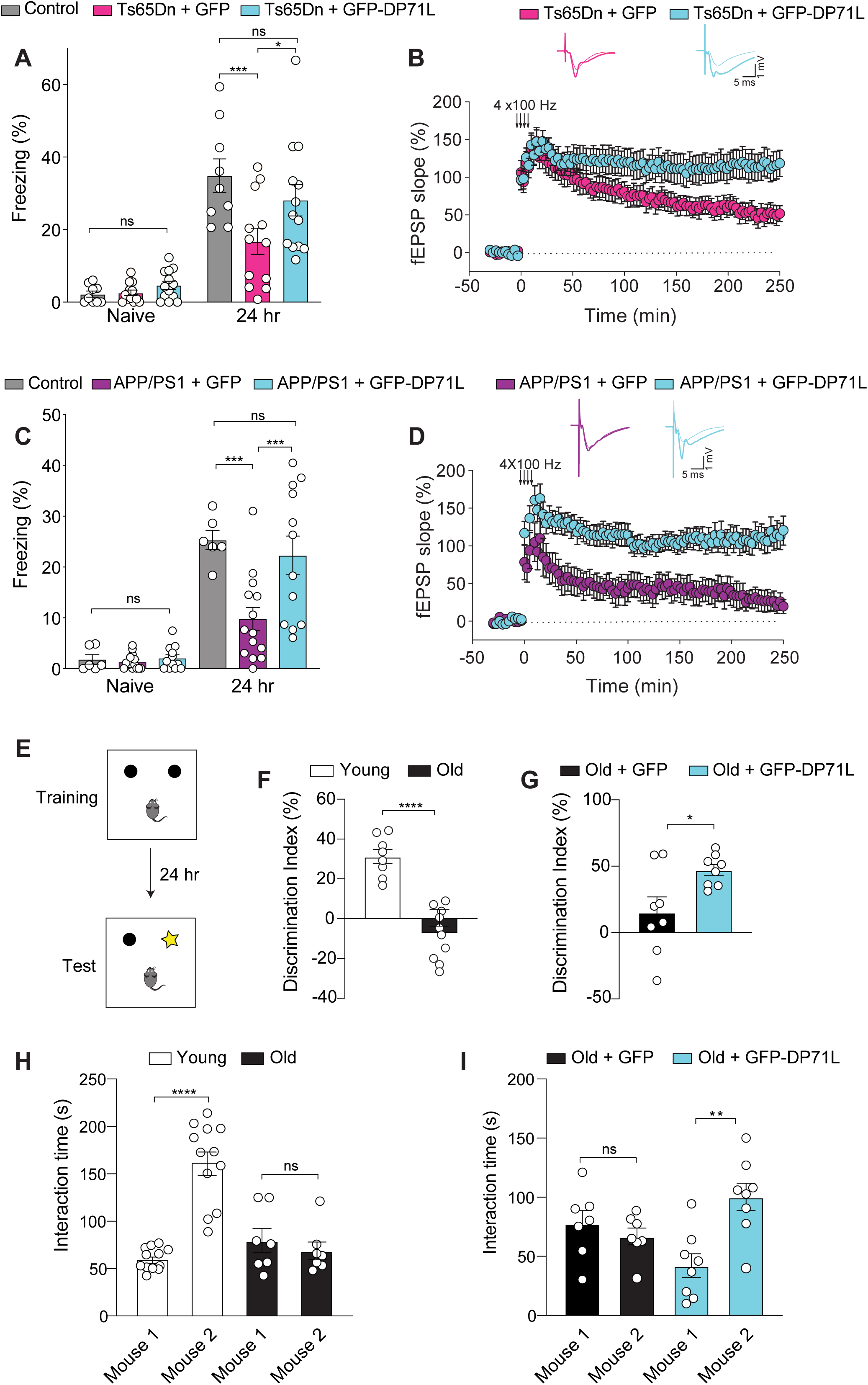
A single injection of DP71L restores long-term synaptic plasticity and memory across various cognitive disorders. (B) Long-term contextual fear memory in WT control mice (*n* = 9) and Ts65Dn mice injected with either AAV-GFP (*n* = 12) or AAV-GFP-DP71L (*n* = 13) [*F*2,62 = 4.18; Control vs. Ts65Dn+GFP: *t* = 3.99, *P* < 0.001; Ts65Dn+GFP vs. Ts65Dn+DP71L: *t* = 2.76, *P* < 0.05; Control vs. Ts65Dn+DP71L: *t* = 1.51, *P* = 0.41]. Freezing behavior was measured before (naïve) and 24 hours after training. (C) L-LTP induced by four trains of high frequency stimulation (HFS, 4 x 100 Hz) in Ts65Dn+GFP mice (*n* = 10) or Ts65Dn+GFP-DP71L mice (*n* = 8) (at 250 min: *U* = 13, *t* = 2.716, *P* < 0.05). (D) Long-term contextual fear memory in control mice (*n* = 6) and APP/PS1 mice injected with either AAV-GFP (n = 14) or AAV-GFP-DP71L (n = 12) [*F*2,58 = 6.43, Control vs. APP/PS1+GFP: *t* = 4.42, *P* < 0.001; APP/PS1+GFP vs. APP/PS1+DP71L: *t* = 4.37, *P* < 0.001; Control vs. APP/PS1+DP71L: *t* = 0.88, *P* > 0.99]. (E) L-LTP induced by four trains of high frequency stimulation (HFS, 4 x 100 Hz) in APP/PS1+GFP (*n* = 10) and APP/PS1+GFP-DP71L (*n* = 10) mice (*t* = 3.92, *P* < 0.001). (F) Schematic of the novel object recognition paradigm. (G) Novel object discrimination index 24 hours after training in Young (*n* = 8) and Old (*n* = 10) mice (*t* = 6.96, *P* < 0.0001). (H) Novel object discrimination index 24 hours after training in Old+GFP (n = 8) and OId-GFP+DP71L (n = 8) mice (*t* = 2.58, *P* < 0.05). (I) Three chamber social novelty in Young (*n* = 12) and Old (n = 7) mice (*F*1,34 = 30.20, Young: *t* = 8.22, *P* < 0.0001; Old: *t* = 0.64, *P* > 0.99). Three chamber social novelty in Old+GFP (*n* = 7) and Old+GFP-DP71L (n = 8) (*F*1.26 = 11.42, Old- GFP: *t* = 0.73, *P* > 0.99; Old+GFP-DP71L: *t* = 4.16, *P* < 0.01). Data are mean ± SEM. \**P* < 0.05, \*\**P* < 0.01, \*\*\**P* < 0.001, \*\*\*\**P* < 0.0001).

Given that Alzheimer’s disease frequently develops in individuals with Down syndrome by age 45^53^ and that the associated cognitive decline is causally linked to activation of the ISR^54–56^, we next tested the efficacy of DP71L in reversing the long-term memory deficits associated with Alzheimer’s disease. To this end, we used the widely employed APP/PS1 mouse model, which expresses mutant forms of human amyloid precursor protein (APP) and presenilin 1 (PS1), both of which are associated with familial Alzheimer’s disease (AD)^57^. These transgenic mice develop progressive cognitive deficits, including impairments in learning and memory. As expected, we found that GFP-DP71L reduced the ISR in the brains of APP/PS1 mice (**Figure S11B**). More importantly, we observed that DP71L- treatment reversed the deficits in long-term memory and long-lasting synaptic plasticity in APP/PS1 mice (**Figure 7C-D**).

We next explored the potential role of DP71L in rejuvenating brain function and reversing age-related cognitive decline, a process that has been associated with the activation of the ISR^16,58^. While long-term fear memory was similar in young and old mice (**Figure S12A**), we found that old mice exhibit impaired long-term object recognition memory, as they could not distinguish between familiar and novel objects (**Figure 7E-F**). Notably, injection of GFP-DP71L into the brain of aged mice not only inhibited the ISR (**Figure S11C**), but also restored their deficits in long-term object recognition memory (**Figure 7G**).

To evaluate whether DP71L could mitigate other behavioral abnormalities associated with aging, we focused on social behavior, as social deficits have been reported to precede the cognitive deficits during aging^59^. When we tested social behavior using the 3-chamber sociability test, which requires the mice to distinguish between an empty cup and a cup with a mouse (**Figure S12B**), we found young and old mice showed similar sociability (**Figure S12C**) and DP71L-treatment did not alter it (**Figure S12D**). However, in the social novelty test, young mice were able to distinguish between a familiar and new mouse, whereas old mice did not, indicating impaired social behavior (**Figure 7H**). Importantly, DP71L-treatment rescued the deficits in social behavior in aged mice (**Figure 7I**). Taken together, these data strongly support the potential for DP71L as a gene therapy for cognitive disorders of diverse origins, including age-related cognitive decline.

Finally, we wondered whether a molecule like DP71L that reverses cognitive decline in several pathological conditions could also potentially enhance long-term memory in healthy, non-diseased mice. Typically, both humans and mice generally require strong and repeated training regimens to form long-term memories, as weak training paradigms usually do not elicit robust long-term memory formation^59^. Consistent with this idea, a weak training paradigm (a single pairing of a tone with a 0.35 mA foot shock) induced only a modest long-term memory in control mice (**Figure S13B**). In contrast, the same training protocol elicited significantly higher long-term memory in DP71L-treated mice, which exhibited reduced ISR in the brain (**Figure S13A-B**). Furthermore, we found that a single stimulus train of 100 Hz elicited a short-lived early-LTP response in control mice, while it produced a sustained and long-lasting potentiation in DP71L-treated mice (**Figure S13C**). Notably, control mice required a strong protocol (four trains of 100 Hz) to achieve a similar long-lasting LTP response to the one observed in DP71L-treated mice with a weak protocol (one train of 100 Hz) (see **Figure S13D**). Hence, these results indicate that DP71L can convert a transient short-lasting mnemonic process into a more long-lasting one.

In conclusion, we found that DP71L not only reverses the long-term memory deficits in a wide range of mouse models of cognitive disorders but also enhances long-term memory in healthy mice, highlighting its potential broader applicability as a therapeutic agent.

## Discussion

Our findings highlight the crucial role the ISR plays in cognitive dysfunction. We discovered that the PPP1R15B^R658C^ variant destabilizes the PPP1R15B•PP1 phosphatase complex, causing a persistent activation of the ISR, which in turn reduces protein synthesis, and leads to impairments in synaptic function and long-term memory (**Figure 1, 2A-C**). More importantly, we provide direct evidence that suppression of the ISR rescued the long-term memory and synaptic deficits in *Ppp1r15b*^R658C^ mice (**Figure 2D-F**), demonstrating that the cognitive deficits associated with the PPP1R15B^R658C^ variant are indeed a direct consequence of persistent ISR activation. Moreover, we found that the increased GABAergic, but not glutamatergic, synaptic transmission in *Ppp1r15b*^R658C^ mice is also the result of persistent ISR activation (**Figure S3**), underscoring the differential susceptibility of distinct neuronal types to persistent ISR activation. These results are consistent with previous findings that ISR inhibition specifically reduces inhibitory synaptic transmission^13,52^ and that reducing the ISR selectively rescues the enhanced inhibitory synaptic transmission and cognitive deficits in a mouse model of Down syndrome^52^.

Interestingly, despite the fact that cell-type specific inhibition of the ISR in glutamatergic and some types of GABAergic neurons facilitates long-term memory formation^18^, the gene expression program induced by persistent ISR activation differs between GABAergic and glutamatergic neurons, with ATF4 being functionally important only in GABAergic neurons (accompanying manuscript, Törkenczy *et al.*). These data support the notion that the ISR impacts its downstream targets through distinct mechanisms in different brain cells.

At the molecular level, the dysregulation of translational efficiency across various mRNAs in *Ppp1r15b*^R658C^ mice indicates that persistent ISR activation reprograms the gene expression landscape in a manner that is different from acute ISR activation (**Figure 3**). A notable example is the absence of feedback regulation typically associated with acute ISR responses (e.g., expression of PPP1R15A^6^; **Figure S9**), which raises intriguing questions about the long-term consequences of chronic ISR activation and highlights the need for further research into ISR phosphatases in both health and disease. Furthermore, single-cell RNA-seq analyses of the brain of *Ppp1r15b*^R658C^ mice reveal that GABAergic and glutamatergic neurons exhibited a more prominent ISR activation and rely on distinct ISR effectors for regulating long-term memory (see accompanying paper, Törkenczy *et al.*). These data indicate that the molecular ISR downstream effectors involved in cognitive processes vary across cell types, findings that were unexpected at the outset.

A unifying perspective: DP71L as a buffer for multiple cognitive disorders. Exploiting evolutionary clues, we characterized DP71L—a small viral orthologue of human phosphatase cofactors primarily composed of the PP1- and eIF2-binding motifs —as a potent pan-ISR inhibitor (**Figure 4**). Its simplified architecture, lacking additional functional N-terminal domains found in larger mammalian counterparts like PPP1R15B and PPP1R15A^24,43^, enables DP71L to effectively engage with the phosphatase PP1. Specifically, mutagenesis and structural studies identified the linker region of DP71L, which connects the PP1 and eIF2 binding motifs, as crucial for its increased binding affinity to PP1 compared to mammalian homologs (**Figure 5-6** and **Figure S14**). This enhanced interaction enables DP71L to inhibit the ISR more effectively. The evolutionary simplicity of DP71L emphasizes the efficiency of small viral genomes, which likely enables viruses to quickly evolve strategies for overcoming cellular stress. Remarkably, we found that DP71L not only alleviates long-term synaptic plasticity and behavioral deficits associated with Down syndrome, Alzheimer’s disease and aging, but it also promotes synaptic function and long-term memory in healthy mice (**Figure 7** and **Figure S12-13**). Hence, DP71L rejuvenates the brain by enhancing the plasticity that underlies long-term memory formation, highlighting its significant therapeutic potential and offering an innovative strategy to address cognitive dysfunction.

How does DP71L reverse the cognitive decline across diverse diseases of multiple origin? We propose that DP71L resets the ISR, thereby buffering cognitive dysfunction by enhancing neural plasticity. Critical periods in postnatal development are crucial windows during which the behavior across many vertebrate species is profoundly shaped^60^. During these phases, neural plasticity peaks, enabling fledgling birds to imprint on their mothers, and human infants to rapidly learn language. However, once these critical periods conclude, the brain’s capacity for learning declines significantly. The ISR has evolved to facilitate the type of neural plasticity that is required for new learning. In birds, auditory imprinting is associated with modulation of the ISR. Specifically, during auditory imprinting the ISR is inhibited, and its activation prevents this crucial process^14^. Remarkably, once the critical period of auditory imprinting has closed, inhibition of the ISR renders the brain more plastic, effectively reopening the critical window of learning. Similar patterns are observed in mammals. Inhibiting the ISR can effectively convert adult mice into a state resembling adolescence regarding their susceptibility to drug-induced changes in synaptic strength and behavior^61–63^. Moreover, the ISR bidirectionally regulates long-lasting synaptic plasticity and ISR inhibition promotes long-term memory formation^12,13,64^.

In multiple cognitive disorders the ISR is persistently activated^1^. Despite different underlying etiologies, all these pathological conditions share a common feature: disruption of protein homeostasis that leads to activation of the ISR^1^. Interestingly, in the accompanying manuscript (Törkenczy *et al.*), based on single cell analysis, we defined a signature of persistent ISR activation indicative of cognitive impairment. Persistent ISR activation hampers neuronal plasticity thus depleting the brain’s buffer capacity, resulting in a significant decline in long-term plasticity and long-term memory formation.

Notably, we found that DP71L-mediated inhibition of the ISR buffers the cognitive decline associated with a wide range of cognitive disorders. Thus, the ISR serves as a “rheostat” finely tuning plasticity and the buffering capacity essential for effective mnemonic processing.

Finally, in this work, we demonstrate the power of intersecting genetics, electrophysiology, behavior, molecular biology, structural biology, evolutionary adaptation, and therapeutic innovation. Our initial studies on a phosphatase cofactor variant associated with human disease (PPP1R15B^R658C^) led us to define the unique architecture and robust function of the viral phosphatase cofactor (DP71L) in ISR modulation, positioning it as a powerful intervention for reversing cognitive impairments across various pathological contexts. Notably, to date, there are approximately 391 known proteins across diverse kingdoms of life that are orthologous to DP71L, representing a largely untapped reservoir for developing innovative treatments for neurological disorders.

## Methods

### Mouse husbandry

*Ppp1r15b*^R658C^ mice were generated using CRISPR/Cas9 editing at the Genetically Engineered Mouse Core Facility at Baylor College of Medicine, as previously described^65^. In brief, embryos were microinjected with donor DNA containing the R658C mutation, Cas9 mRNA (Sigma, St. Louis, MO), and gRNA from Synthego (Redwood City, CA). The gRNA sequence 5’- AGTGGTGATGAGGATCGCAA-3’ was used in combination with a donor sequence containing the R658C mutation and a silent TC to CT mutation at the cut site, which also introduced a DpnII restriction site for downstream genotyping. Injected embryos were transplanted into surrogate dams and genome editing confirmed with Sanger sequencing after amplification with the primers 5’- GGATAACAAACAAACAGAGAAACACAGAGTTCC-3’ and 5’-GGAAACTCTCAAGTAAGAGACAGA GTGG-3. Potential off-target gene editing was analyzed by Sanger sequencing of the top 20 most likely off-target loci predicted from the COSMID analysis tool as previously described^66^. For colony maintenance DNA was prepared by isolating ear tissue with protease K in polymerase buffer overnight at 52°C. Insoluble material was then sedimented and the supernatant was used for PCR amplification with the primers above. The PCR product was then digested with DpnII for 1 h prior to analysis with gel electrophoresis. R658C mutant DNA lacks the restriction site and is undigested in contrast to WT DNA. *Ppp1r15b*^R658C^*-Eif2a1^S/A^* mice were generated by crossing *Ppp1r15b*^R658C^ mice with *Eif2a1^S/A^* mice, which were previously reported^12,67^. APPswe/PSEN1dE9 (stock #<u>004462</u>) and Ts65Dn mice (stock #005252) were obtained from Jackson Labs (Bar Harbor, ME) and were genotyped as recommended by Jackson Labs. Experiments were conducted on male and female mice aged 10-16 weeks old for all mouse lines except APPswe/PSEN1dE9, which were 32-40 weeks of age. Aged mice (18-20 months old) were obtained from Jackson Labs. Animals were kept under a 12h / 12h light/dark cycle (lights on at 7:00 am) and had access to food and water *ad libitum*. Animal care and use procedures were approved by Baylor College of Medicine’s and Charles River Laboratories’ Animal Care and Use Committees in accordance with guidelines established by the National Institutes of Health.

### Lentivirus and AAV

GFP, GFP-DP71L, and GFP-ΔPPP1R15B were cloned into pLVX to generate the transfer plasmids for lentiviral transduction. To generate lentivirus, 293T cells were cotransfected with transfer plasmid, psPAX2, and pMD2.G at a 1:1: 0.5 ratio with lipofectamine 3000. Cell culture medium was replaced after 16h and then harvested at 24, 48, and 72 h thereafter, and pooled. Supernatant was filtered over a 0.45μM PES membrane. Virus was diluted and applied to cells with 10μg/mL polybrene to infect HEK293T cells.

DP71L was cloned into an AAV2 pCAG-GFP vector to generate an n-terminal tag on DP71L. For functional experiments to assess eIF2-P, a 2A protease peptide separated GFP from DP71L. For AAV production, we used either the Neuroconnectivity Core facility at Baylor College of Medicine (NCC) or the University of Pennsylvania’s Gene Therapy Program Vector Core (GTP). The NCC and GTP core facilities used the DJ8 serotype for viral production and quantified the virus concentration after purification using a qPCR-based approach.

<u>AAV injection:</u> Mice were anesthetized in accordance with institutional procedures using isoflurane (2- 3%), placed on a Leica stereotactic frame (Buffalo Grove, IL). AAV (∼10^11^ copies/mL for GFP alone and∼10^12^ copies/mL for DP71L virus; 1μl) and injected bilaterally with a 20 μl glass syringe attached to a motorized KD Scientific syringe pump (Holliston, MA) in the hippocampus (Bregma: -2.30 AP, ±1.80 ML, -1.40 DV (CA1) and -1.94 AP, ±2.20 ML, -2.20 DV (CA3) at a rate of 0.1 μl/min. Following the infusion, the injector remained for 10 min to allow complete diffusion of the solution from the tip of the injector.

### Cell culture

HEK293T cells stably expressing DP71L-GFP or ΔPPP1R15B-GFP were generated and maintained in DMEM without pyruvate supplemented with FBS, glutamine and HEPES, pH7.5. Cells were treated with either oligomycin (1 μM, Sigma, St. Louis, MO)), neratinib (1 μM, Sigma, St. Louis, MO), or thapsigargin (100 nM, Sigma, St. Louis, MO) for 3 hours at 37° C. For poly(I:C) experiments, cells were transfected with 285 ng poly(I:C) from Invitrogen (San Diego, CA) per 1 cm^2^ using Lipofectamine 2000 (Thermo Fisher, Waltham, MA) and 6 hours post-transfection, they were washed with ice-cold PBS and harvested in lysis buffer.

### Protein expression and purification

DP71L protein (with N-term 6xHis-SUMO and C-term MBP tags) and untagged human PP1A (residues 7-300 with catalytically dead D64A mutation) were cloned into a single pETDuet-1 vector (Novagen, Reno, NV). The plasmid was transformed into chemically competent T7 Express *lys*Y/I^q^ cells (New England Biolabs, Ipswich, MA). *E. coli* was expanded in 2xYT media (with 1 mM MnCl2), and expression was induced with IPTG added to 0.5 mM. Following 18 hours of expression at 22° C, the cells were harvested by centrifugation and washed with PBS. Cell pellets were then resuspended in cold lysis buffer containing 20 mM Tris-HCl pH 7.5, 500 mM NaCl, 1 mM MnCl2, 20 mM imidazole, 1 tablet/50 mL protease inhibitor cocktail (Thermo Fisher Scientific, Waltham, MA), and 1 mM PMSF. Cells were lysed through four cycles of high-pressure homogenization using the EmulsiFlex-C3 (Avestin, Ottawa, ON, Canada) then clarified by centrifugation for 1 hour at 18,000 x g at 4° C. All subsequent purification was carried out using an AKTA Pure system (Cytiva, Marlborough, MA) at 4° C. The lysate was vacuum filtered and loaded onto a HisTrap FF column (Cytiva, Marlborough, MA) and equilibrated following the manufacturer’s instructions. The column was washed with 10 column volumes (CV) of 90% HisTrap buffer A (20 mM Tris-HCl pH 7.5, 500 mM NaCl, 1 mM MnCl2)/10% HisTrap buffer B (20 mM Tris-HCl pH 7.5, 500 mM NaCl, 1 mM MnCl2, 200 mM imidazole). Bound protein was eluted using a 10-100% gradient of HisTrap buffer B over 10 CV. Peak fractions were pooled and the 6xHis-SUMO tag was cleaved overnight using 6xHis-Ulp1 protease while dialyzing against cleavage buffer (20 mM HEPES-KOH pH 7.5, 500 mM KCl, 1 mM MnCl2). Cleaved protein was passed over an equilibrated gravity column containing HisPure NiNTA resin (Thermo Fisher Scientific, Waltham, MA) and the flow-through was collected and concentrated. The concentrated sample was purified further by size exclusion chromatography using a HiLoad 16/600 Superdex200 pg column (Cytiva, Marlborough, MA) equilibrated in SEC buffer (20 mM HEPES-KOH pH 7.5, 150 mM KCl, 1 mM MnCl2). The purified dimeric complex was flash frozen and stored at -80°C until use.

G-actin from rabbit muscle (Cytoskeleton, Denver, CO) was resuspended to a final concentration of 5 mg/mL in G-actin buffer (5 mM Tris–HCl pH 8, 0.2 mM ATP, 0.5 mM DTT, 0.2 mM CaCl2) and treated with a 10x molar excess of Latunculin B (Abcam, Cambridge, UK) overnight at 4° C while rotating. DNAseI, grade II from Bovine pancreas (MilliporeSigma, Burlington, MA), was resuspended to 10 mg/mL in DNAseI buffer (10 mM Tris–HCl pH 7.5, 50 mM NaCl, 1 mM CaCl2) and added to the G-actin mixture at a 1.5x molar excess then incubated for 2 hours while mixing at 4° C. The combined sample was then purified by size exclusion chromatography using a Superdex75 Increase 10/300 GL column (Cytiva, Marlborough, MA) equilibrated with G-actin SEC buffer (10 mM HEPES-KOH pH 7.5, 150 mM KCl and 1 mM CaCl2, 0.2 mM ATP and 0.2 mM TCEP). Peak fractions containing dimeric DNaseI/G-actin were pooled, concentrated, and flash frozen prior to storage at -80° C.

Human eIF2α (residues 2-187) with an N-terminal 6xHis tag was expressed in NiCo21 (DE3) cells (NEB #C2529) grown in of ZYP-5052 autoinduction media for 18 hours at 20° C. Following expression, cells were harvested by centrifugation then resuspended with 3 mL lysis buffer per gram of wet pellet (20 mM Tris HCl pH 7.5, 500 mM NaCl, 20 mM imidazole, 1 tablet/50 mL EDTA-free protease inhibitor cocktail (Thermo Fisher Scientific, Waltham, MA), 1 mM PMSF per gram of wet pellet and lysed by sonication using a Branson sonifier SFX 550 (10s on/off pulses for 10 minutes at 60% power). The lysate was cleared by spinning at 18000 x g for 30 minutes at 4 °C. All subsequent purification was carried out using an AKTA Pure system (Cytiva, Marlborough, MA) at 4° C. The lysate was vacuum filtered and loaded onto a HisTrap FF column (Cytiva, Marlborough, MA) equilibrated following the manufacturer’s instructions. The column was washed with 10 CV of 92% HisTrap buffer A (20 mM Tris-HCl pH 7.5, 500 mM NaCl), 8% HisTrap buffer B (20 mM Tris-HCl pH 7.5, 500 mM NaCl, 250 mM imidazole) and with a further 5 CV using 25% buffer B. Bound protein was eluted in 100% HisTrap buffer B until the elution peak diminished. Elution fractions were pooled and MgCl2 and ATP were added to a final concentration of 5 mM and 2.5 mM, respectively, followed by addition of poly (I:C) activated PKR^65^ at a ratio of 1:50 (w/w) for *in vitro* phosphorylation. The reaction was carried out for 1 h at 37° C then subjected to size exclusion chromatography using a Superdex75 Increase 10/300 GL column (Cytiva, Marlborough, MA) to remove any PKR. Completion of the phosphorylation reaction was confirmed by SDS PAGE using a Phos-Tag gel (Fujifilm-Wako, Richmond, VA). The final sample was concentrated, flash frozen in liquid nitrogen, and stored at -80° C.

Cryo-EM sample preparation and data collection:

The DP71L-PP1A-G-actin-DNAseI complex was assembled by combining 40 nmol of the PP1A/DP71L heterodimer, eIF2-P (residues 2-187), and the G-actin/DNAseI dimer and purifying the sample by size exclusion chromatography using a Superdex200 Increase 10/300 GL column (Cytiva, Marlborough, MA). Peak fractions were analyzed by SDS-PAGE and fractions containing all the components were pooled and concentrated to 5.5 mg/mL.

Quantifoil (Au) 1.2/1.3 25 nm, 300 mesh grids were glow discharged using a Pelco easiGlow for 30 s at 15 mA used immediately for plunge freezing using a Vitrobot Mark IV (Thermo Fisher Scientific, Waltham, MA). 3.5 µL of sample was applied to a grid, excess liquid was blotted away for 1.5 s with a blot force of -5 at 4.4 °C and 100% humidity followed by plunge freezing in 100% liquid ethane. All grid screening and data collection was done at the Altos Labs Electron Microscopy Hub, using a Krios G4 TEM operating at 300 kV equipped with a Falcon4i detector and a SelectrisX energy filter with a slit width of 10 eV (Thermo Fisher Scientific, Waltham, MA). 5,236 exposures were collected with a total dose of 50 electrons/Å^2^ using a parallel beam at a flux rate of ∼ 7 electrons/pixel/sec set to a calibrated physical pixel size of 0.9286 Å and a defocus range of -0.8 to -1.8 µm using Aberration Free Image Shift (AFIS) between acquisitions. Data collection was automated using EPU software version 3.6.0 (Thermo Fisher Scientific, Waltham, MA) and recorded at the internal frame rate of the camera in counting mode and written in EER format. Motion correction and early image processing was executed in cryoSPARC Live version 4.4.0^66^. EER frames were combined into 50 fractions each representing 1 electron/Å^2^ exposure and processed without up sampling. Objective lens and coma astigmatism were corrected prior to data acquisition.

### Cryo-EM image analysis, model building, and structure prediction

All data processing was carried out using cryoSPARC v4.4+ unless otherwise stated. An initial subset of particles was picked using a blob picker set to 80-120 Å particle diameter range and subjected to 2D classification to create templates for template-based picking. 1,315,944 particles were picked using the template picker set to 128 Å particle diameter and extracted with a box size of 320 pixels, Fourier cropped to 160 pixels for initial processing. The particles were used for *ab initio* reconstruction of three classes which were refined by heterogeneous refinement using all particles with the refinement box size set to 192 voxels, resulting in two classes containing 780,355 particles which were uncropped and used for further refinements. Two iterative rounds of 2D and 3D classification were used to clean the particle stack, removing damaged and partially assembled complexes. The resulting stack of 309,745 particles was used for homogenous refinement with an initial low-pass filter of 16 Å, minimizing over per-particle scales, and adaptive marginalization over poses and shifts, yielding a map with an overall resolution of 3.22 Å. The refined particles and volume were next used for reference-based motion correction to estimate per-particle motion and movie frame dose weighting. The motion corrected particles were used for a final homogenous refinement of the previous map (low-pass filtered to 16 Å), with per-particle defocus refinement, per-group CTF refinement, and tilt, trefoil, tetrafoil, spherical aberration, and anisotropic magnification fitting enabled. Overall resolution of the final map was 3.1 Å as measured at the FSC=0.143 criterion after FSC-mask auto-tightening. To improve the map quality around individual subunits, local masks were generated surrounding G-actin, DNAseI, and PP1-DP71L-eIF2α, respectively, and used for local refinement. Alignment parameters for local refinement included enabling using poses and shift Gaussian priors (5° and 3 Å standard deviation, respectively) and enabling re-centering poses and shifts each iteration. The locally refined maps were used for final model building.

### Model building and structure prediction

All model building and refinement were done using ISOLDE^70^ within UCSF ChimeraX v1.7.1 or v1.8.0^45^. AlphaFold2-multimer^71^ was used to generate a starting model containing human PP1A (residues 7-300), ASFV DP71L, human eIF2α (residues 2-187), bovine DNAseI, and rabbit G-actin. The model was fit into the cryo-EM map and first globally relaxed by running the simulation on all chains with default settings. Each residue of PP1A and DP71L was then manually inspected and fit into the density with the ISOLDE molecular dynamics simulation enabled. Lastly, molecules not modeled by AlphaFold, such as an ATP molecule bound to G-actin and Mn^2+^ ions were added to the model based on previous knowledge and observed cryo-EM density. A final refinement of the model was carried out against half-maps using Servalcat^72,73^ and the resulting statistics are reported in **Table IV**. Experimental structures are deposited to the wwPDB with accession codes EMD-49124, EMD-49162, EMD-49163, EMD-49164, EMD-49223, PDB-9NB9.

Structures of ΔPPP1R15B (PPP1R15B residues 638-697aa) and DP71L were predicted using both AlphaFold2-Multimer implementation within the ColabFold notebook server^74^ and OpenFold^49^. Structures were predicted using the pdb100 template mode, and the top ranked structure was relaxed using Amber with all other settings kept to default. All figures which include experimental and predicted structures were generated using UCSF ChimeraX^45^.

### Immunoprecipitation (IP) and Western blotting

IP and western blotting were performed as previously described^52,75^, with some modifications, Briefly, constructs for IP were generated without the P2A sites in the AAV2 pCAG backbone to allow expression of GFP-tagged proteins. 293T cells were transfected with either the vector alone or AAV2 pCAG constructs lacking P2A sites using lipofectamine 3000. After 16 hours, cells were washed with ice-cold PBS and harvested. Lysates were prepared as above except in IP lysis buffer: 40 mM NaCl, 50 mM HEPES, 2 mM EDTA, 1mM Na3VO4, 50 mM NaF, 10mM glycerophosphate, 0.3% CHAPS, and protease inhibitor tablet (ROCHE, Basel, Switzerland). 1mg of total protein was used to immunoprecipitate GFP-tagged protein by incubating overnight at 4° C with tumbling with GFP-trap magnetic resin (Proteintech Group, Rosemont, IL). The supernatant was removed prior to washing in IP lysis buffer four times. Immunocomplexes were eluted using 1X Laemmli buffer and resolved with SDS-PAGE prior to western blotting.

For Western blotting, tissue lysates were prepared in the following lysis buffer: 50mM NaCl, 200mM HEPES pH7.5, 1mM EDTA, 10% glycerol, 25mM Δ-glycerophosphate, 50mM NaF and 2mM Na3VO4 and 1X protease inhibitor (ROCHE, Basel, Switzerland). Insoluble material was removed and the protein concentration in the supernatant was quantified using Bradford analysis (Biorad, Hercules, CA), prior to SDS-PAGE. Antibodies used in this study: 119A11 eIF2-P (Cell Signaling Technology, Danvers, MA) or E90 ab32157 eIF2-P (Abcam, Cambridge, UK), D7D3 XP eIF2-T (Cell Signaling Technology, Dnvers, MA), M0802 GFP (Abiocode, Agoura Hills, CA), or A11192 GFP (Thermo Fisher Scientific, Waltham, MA), MABE343 puromycin (MilliporeSigma, Burlington, MA), sc7482 clone E9 PP1α (Santa Cruz Biotechnology, Santa Cruz, CA), PA5-31177 eIF2α (Thermo Fisher Scientific, Waltham, MA), A301-742A eIF2α (Bethyl Laboratories, Montgomery, TX), A300-721A eIF2α (Bethyl Laboratories, Montgomery, TX), 10449-1-AP PPP1R15A (Proteintech Group, Rosemont, IL), 14634-1- AP PPP1R15B (Proteintech Group, Rosemont, IL), 60004-1-IG GAPDH (Proteintech Group, Rosemont, IL).

### qRT-PCR

Total RNA was extracted from brain tissue by first homogenizing in TRIzol followed by RNA cleanup using the Zymo RNA clean and concentrator kit (Irvine, CA). RNA was quantified and cDNA was generated using SuperScript VILO (Thermo Fisher Scientific, Waltham, MA). qPCR was performed using PowerUp SYBR Green Master Mix, according to the manufacturer’s protocol (Applied Biosystems, Carlsbad, USA) to quantify relative expression levels of the ATF4 target genes. Expression of the mouse GAPDH was used to normalize each gene expression levels. Primer sequences are provided in **Table III**.

### Polysome profiling

Polysome profiling was conducted as described previously^52^, with some modifications. Fresh 12 mL sucrose density gradients (10–50%) were prepared using 10 mM HEPES-KOH (pH 7.6), 5 mM MgCl2, and 150 mM KCl, supplemented with 200 U/mL RNasin Rnase inhibitor (Promega, Madison, WI). The gradients were maintained at 4°C for at least 2 hours prior to use. Mouse brain tissue was dissected in a cutting solution composed of 1X HBSS, 2.5 mM HEPES-KOH (pH 7.6), 35 mM glucose, 4 mM NaHCO3, and 100 µg/mL cycloheximide (Sigma-Aldrich, St. Louis, MO). After dissection, the tissue was washed in ice-cold PBS containing 100 µg/mL cycloheximide, followed by centrifugation at 3000 rpm for 10 minutes at 4° C. Tissue, was then lysed in lysis buffer [10 mM HEPES-KOH (pH 7.4), 5 mM MgCl2, 150 mM KCl, 0.5 mM DTT, 100 U/mL RNasin Rnase inhibitor (Promega, Madison, WI), 100 µg/mL cycloheximide, and EDTA-free protease inhibitors (Roche, Indianapolis, IN)] and centrifuged at 2000 x g for 10 minutes at 4° C. The supernatant was transferred to a pre-chilled tube, supplemented with 0.5% NP-40, and incubated on ice for 10 minutes. Subsequent centrifugation was performed at 14,000 rpm for 10 minutes at 4°C.

The resulting supernatant was either layered onto the sucrose gradient or reserved for total RNA isolation. Gradients were centrifuged in a SW-40Ti rotor at 35,000 rpm for 2 hours at 4° C. After centrifugation, fractions were collected by piercing the tube with a Brandel tube piercer and introducing 70% sucrose from the bottom. The eluting material was monitored for absorbance using an ISCO UA-6 UV detector. RNA was extracted from the collected fractions using TRIzol, following the manufacturer’s instructions (Life Technologies, Carlsbad, CA). Experiments were performed in several biological replicates for each group.

### Puromycylation

Protein synthesis was measured using SUnSET, a non-radioactive labeling method to monitor protein synthesis was previously described^52,76^. Briefly, brain slices were cut (300 μm) with a McIlwain Tissue Chopper (Mickle, UK) and incubated for 1 hour at room temperature in oxygenated (95% O2, 5% CO2) ACSF followed by incubation at 32° C for 1 hour in oxygenated ACSF (124 mM NaCl, 2.0 mM KCl, 1.3 MgSO4 mM, 2.5 mM CaCl2, 1.2 mM KH2PO4, 25 mM NaHCO3, and 10 mM glucose) prior to treatment as we previously described^52^. Puromycin (10 μg/μl, dissolved in oxygenated ACSF) was bath applied to the slices for 20 min followed by an ACSF wash without puromycin. Tissue slices were then snap-frozen on dry ice and stored at -80° C until use. Frozen slices were lysed in WB lysis buffer. Puromycin incorporation was detected by Western blot as we previously described^52^. The density of the resulting bands was quantified using ImageJ.

### Ternary complex analysis

For immunoprecipitation, antibody-conjugated beads were prepared from Dynabeads M270 Epoxy beads using antibody directed against eIF2α, as previously reported^77^. Brain tissue was lysed in IP Lysis Buffer (150mM NaCl; 50mM Tris-HCl, pH 8.0, 1% Triton X-100, 100mM sodium fluoride, 50mM β-glycerophosphate, and 10mM sodium pyrophosphate), cleared, and then quantified with Bradford reagent. Immunocomplexes were precipitated overnight with antibody-conjugated beads from ∼1mg input material, washed with IP lysis buffer, and recovered by eluting either with Laemmli buffer (without DTT) or with TRIzol. Laemmli-eluted material was subjected to SDS-PAGE. TRIzol-eluted material was used for total RNA isolation and cDNA synthesis prior to PCR with the following primers: Met-tRNAi^Met^ forward 5’- AGCAGAGTGGCGCAGC-3’, and Met- tRNAi^Met^ reverse 5’- TAGCAGAGGATGGTTTCGATCC-3’.

### Ribosome profiling and RNA-seq

Ribosome profiling was conducted as previously described^31,78,79^. Briefly, brain tissue was harvested into polysome buffer and separated into two aliquots, one for ribosome profiling and the other for RNA-seq. The ribosome profiling aliquot was digested at 4° C for 1h with RNaseT1 at 2.5 U/μl. The digestion of ribosome-bound mRNAs was stopped with the addition of SuperaseIn (Thermo Fisher Scientific, Waltham, MA). Digested samples were then loaded onto sucrose gradients, as described above, and RNA from 80S peaks was purified using TRIzol followed by 15% denaturing Urea gels. Gel-purified ribosome protected fragments (RPFs) were cleaned up with a Zymo RNA clean and concentrator prior to library preparation with a NEB small RNA library preparation kit (New England Biolabs, Ipswich, MA). Following cleanup of the final amplified libraries, samples were quantified using a Qubit (Thermo Fisher Scientific, Waltham, MA). Samples were sequenced by the Advanced Technologies Genomics Core at MD Anderson Cancer Center (Houston, TX) using the Nextseq500 platform (Illumina, San Diego, CA).

The corresponding samples designated for RNA-seq were processed as follows. Total RNA was extracted using TRIzol, and RNA from the aqueous phase was purified using the Zymo RNA Clean and Concentrator-25 kit (Irvine, CA). Samples were quantified using the nanodrop spectrometer (Thermo Fisher Scientific, Waltham, MA) and sent for sequencing at Novogene (Sacramento, CA) using the Novoseq platform (Illumina, San Diego, CA).

FASTX-clipper and -barcode splitter (http://hannonlab.cshl.edu/fastx_toolkit/) were used to remove the linker sequences and de-multiplex ribosome profiling data, respectively. Unique molecular identifiers and sample barcodes were subsequently removed using a custom Python script. Reads corresponding to rRNAs were excluded using Bowtie v1.1.2 (http://bowtie-bio.sourceforge.net/) and the remaining reads were aligned to the GRCm39/mm39 transcriptome using tophat v2.1.1 (https://ccb.jhu.edu/software/tophat/) using flag sets as follow: --b2-very-sensitive --transcriptome-only --no- novel-juncs --max-multihits=64 flags. We then used the plastid cs program (Dunn and Weissman, 2016) to calculate counts per gene and normalized counts per gene (in reads per kilobase per million mapped reads, or RPKM), with counts assigned to the ribosome P-site determined by the plastid psite program. RNA-seq data were analyzed similarly. Additional analysis and visualizations were performed in Python 2.7 using plastid, Biopython, Numpy, Pandas, and GSEApy.

Translation efficiency (TE) is defined as the ratio of normalized ribosome footprint RPKM to normalized mRNA RPKM. TE and mRNA fold changes and adjusted p values were calculated from raw counts with DESeq2 ^80^. TE changes were calculated using the design formula ∼sample + sample:assay, where the sample interaction term denotes the sample and the assay interaction term specifies whether counts are derived from RNA-seq or ribosome profiling. mRNA changes were calculated from RNA-seq counts using the design formula ∼“sample”.

### Behavioral experiments

All experiments were conducted during the light cycle and performed and analyzed blind to the genotypes or treatment.

- *Contextual Fear Conditioning*: Experiments were performed as previously described^13,52^,with some modifications. Briefly, mice were first handled for 3 days (for 5-10 min each day) and then habituated to the conditioning chamber for 20 min for another 2 days. On the training day, mice were placed in the conditioning chamber for 2 min (naïve) and then received two-foot shocks (0.75 mA, 2 sec, 90 sec apart), after which the mice remained in the chamber for an additional minute before being returned to their home cages. For weak training mice only received one foot shock (0.35 mA). Twenty-four hours later, mice were re-exposed to the same context for 5 min and “freezing” (immobility except for respiration) responses were recorded using real-time video and analyzed by FreezeView (Actimetrics, Lafayette, IN).
- *Novel Object Recognition:* Object recognition was performed as previously described^81^.

Briefly, mice were handled for 3 days (5-10 min for each day) and then habituated to a black Plexiglas rectangular chamber (31 x 24 cm, height 27 cm) for 10 min under dim ambient light for 5 days. Two identical objects were presented to mice to explore for 5 min, after which, mice were returned to the home cage. Twenty-four hours later, one object was replaced by one novel object and the mouse was again placed in the chamber 5 min. The novel object has the same height and volume but different shape and appearance. Exploration of the objects was defined as sniffing of the objects (with nose contact or head directed to the object) within at 2 cm radius of the objects. Sitting or standing on the objects was not scored as exploration. Behavior was recorded from cameras positioned above the training chamber. Discrimination Index (DI) was computed as DI = (Novel Object Exploration Time – Familiar Object Exploration Time/Total Exploration Time) X 100. To control for odor cues, the open field arena and the objects were thoroughly cleaned with ethanol, dried, and ventilated between mice.

- *Three-Chamber Social Test:* Crawley’s three-chamber test for sociability and social novelty was performed as we previously described^82–84^. Briefly, mice were first acclimatized in an empty 60 × 40 × 23 cm Plexiglas Arena divided into three equally sized interconnected chambers (left, center, right) for a 10 min. During this habituation phase, the mice’s movements were continuously monitored using AnyMaze tracking software, allowing for automated scoring of distance traveled and speed to assess overall activity levels. Following habituation, the sociability of the mice was evaluated during a second 10-minute session. During this period, each mouse had the option to interact either with an empty wire cup (Empty) or a wire cup containing a stranger mouse (Mouse 1), which were age- and sex-matched to the subject mouse. The interaction time was scored by measuring the time the subject mouse spent sniffing (or climbing) upon either the empty cup or the cup containing the stranger mouse. It is noteworthy that the position of the empty cup/stranger mouse in the left or right chamber during the sociability period was counterbalanced between trials to avoid bias.

To evaluate the preference for social novelty, a third 10-minute phase was conducted, wherein a second stranger mouse (Mouse 2) was introduced into the previously empty wire cup. The interaction times with both Mouse 1 and Mouse 2 were again measured, employing the automated AnyMaze system. All interaction time data were scored by trained, independent observers who remained blinded to the treatment conditions throughout the experiment.

### Electrophysiology

Recordings were performed as previously described^13,52^. Briefly, field recordings were performed from CA1 horizontal hippocampal slices (320 μm thick) prepared from mouse brains using a Leica VT 1000S vibratome (Buffalo Grove, IL) at 4° C in artificial cerebrospinal fluid (ACSF). The slices were equilibrated for at least 60 min prior to recording in an interface chamber and continuously perfused (2-3 mL/min) with artificial cerebrospinal fluid ACSF at 28–29° C. Recording electrodes were positioned in the stratum radiatum, where field excitatory postsynaptic potentials (fEPSPs) were recorded with ACSF-filled micropipettes. Stimulation was performed with bipolar stimulating electrodes placed in the CA1 stratum radiatum to activate Schaffer collateral and commissural fibers. The intensity of the 0.1-ms pulses was calibrated to evoke 30-35% of maximal response. A stable baseline of responses at a frequency of 0.033 Hz was established for at least 30 min. Late-LTP was induced by applying four tetanic trains of high-frequency stimulation (100 Hz, 1 s) separated by 5min intervals, while early-LTP was induced by applying one tetanic train of high-frequency stimulation (100 Hz, 1 s).

Whole-cell recordings were performed using a Molecular Devices MultiClamp 700B amplifier (Union City, CA) in a submerged perfusion chamber at 31–32° C using conventional patch-clamp techniques (2-3 mL/min flow), as we previously reported^13,52^. CA1 neurons were visually identified by infrared differential interference contrast video microscopy on the stage of a Zeiss Axioskope F2 upright microscope (Oberkochen, Germany). Patch pipettes (resistances 2 - 5 MW) were filled with patch buffer (140 mM CsCl, 10 mM HEPES, 10 mM Na2-phosphocreatine, 0.2 mM BAPTA, 2 mM ATP, 6 mM MgCl2, 0.2 mM GTP; pH was adjusted to 7.2 and osmolarity to 295∼300 mOsm using a Wescor 5500 vapor pressure osmometer (Logan, UT)). Miniature inhibitory postsynaptic currents (mIPSCs) were recorded at -60 mV in the presence of NBQX (5 μM, 2,3-dihydroxy-6-nitro-7-sulfamoyl-benzo[f]quinoxaline disodium salt), AP5 (25 μM, DL-2-Amino-5-phosphonopentanoic acid) and TTX (1 μM, tetrodotoxin). For recording excitatory currents, CsCl was replaced with 130 mM potassium gluconate and 10 mM KCl. Excitatory postsynaptic currents (EPSCs) were recorded at -70 mV in the presence of 100 μM picrotoxin. All drugs were obtained from Tocris (Ellisville, MO).

### Statistical analysis

Data are presented as mean ± SEM, with individual data points included.

Statistical analyses were performed using the Mann-Whitney U test, Student’s t-test, and one- or two-way ANOVA with Bonferroni post-hoc correction for multiple comparisons. Statistical values of *n*, *t*, *F*, and *P* are provided in the figure legends. A significance level of *P* < 0.05 was used, with \**P* < 0.05, \*\**P* < 0.01, \*\*\**P* < 0.001, and \*\*\*\**P* < 0.0001 indicating increasing levels of significance. GraphPad Prism (La Jolla, CA) software was used for statistical analysis and graphical data representation of molecular, behavioral and electrophysiological experiments.

## Acknowledgements

This work was supported by Altos Labs, Inc., NIMH (M.C.-M.) and the Intramural Research Program of the National Institutes of Health (T.E.D.). We thank Christina Mayer for administrative support, members of Altos Labs’ animal facilities for support, Nimit Jain for discussion about the iMet-tRNA assay. Finally, we also thank members of the M.C.-M. and P.W. labs for comments on the manuscript.

