## Supplementary Material for "Harnessing the Evolution of Proteostasis Networks to Reverse Cognitive Dysfunction"

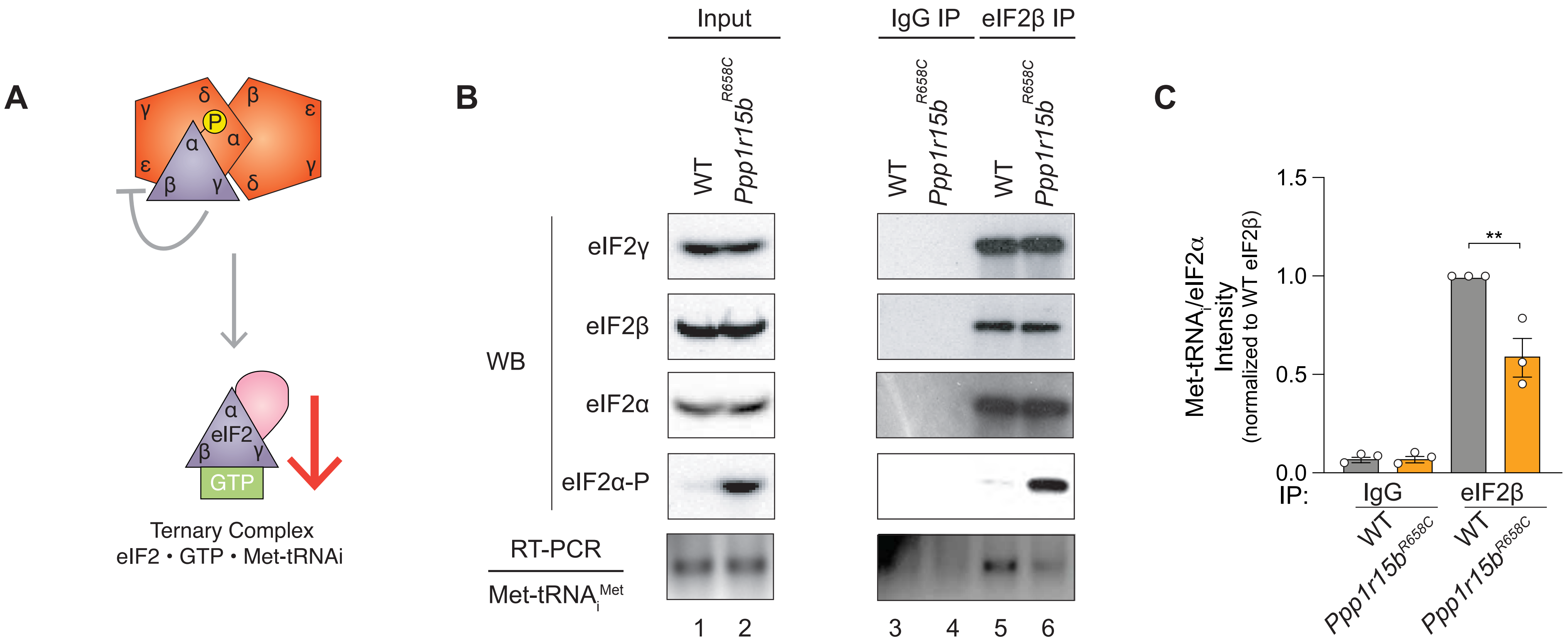


**Fig. S1.** Reineke *et al.*

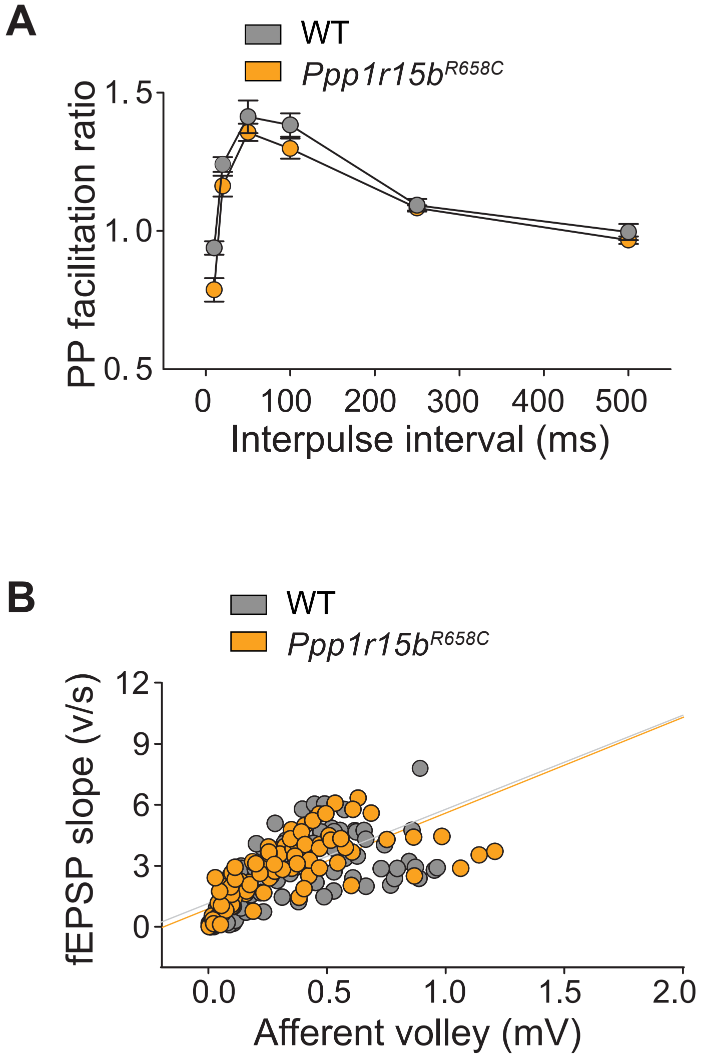


**Fig. S2.** Reineke *et al.*

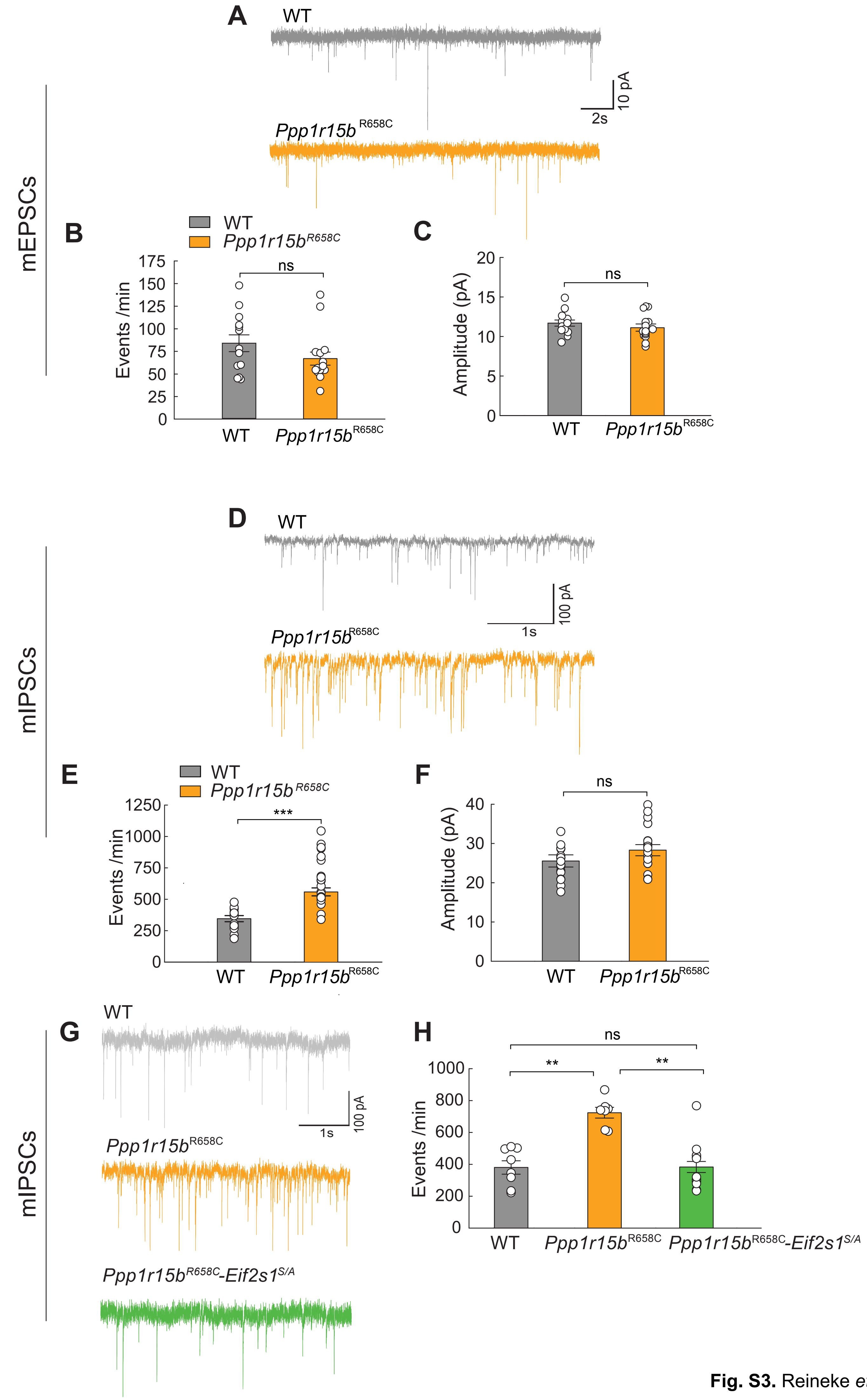


**Fig. S3.** Reineke *et al.*

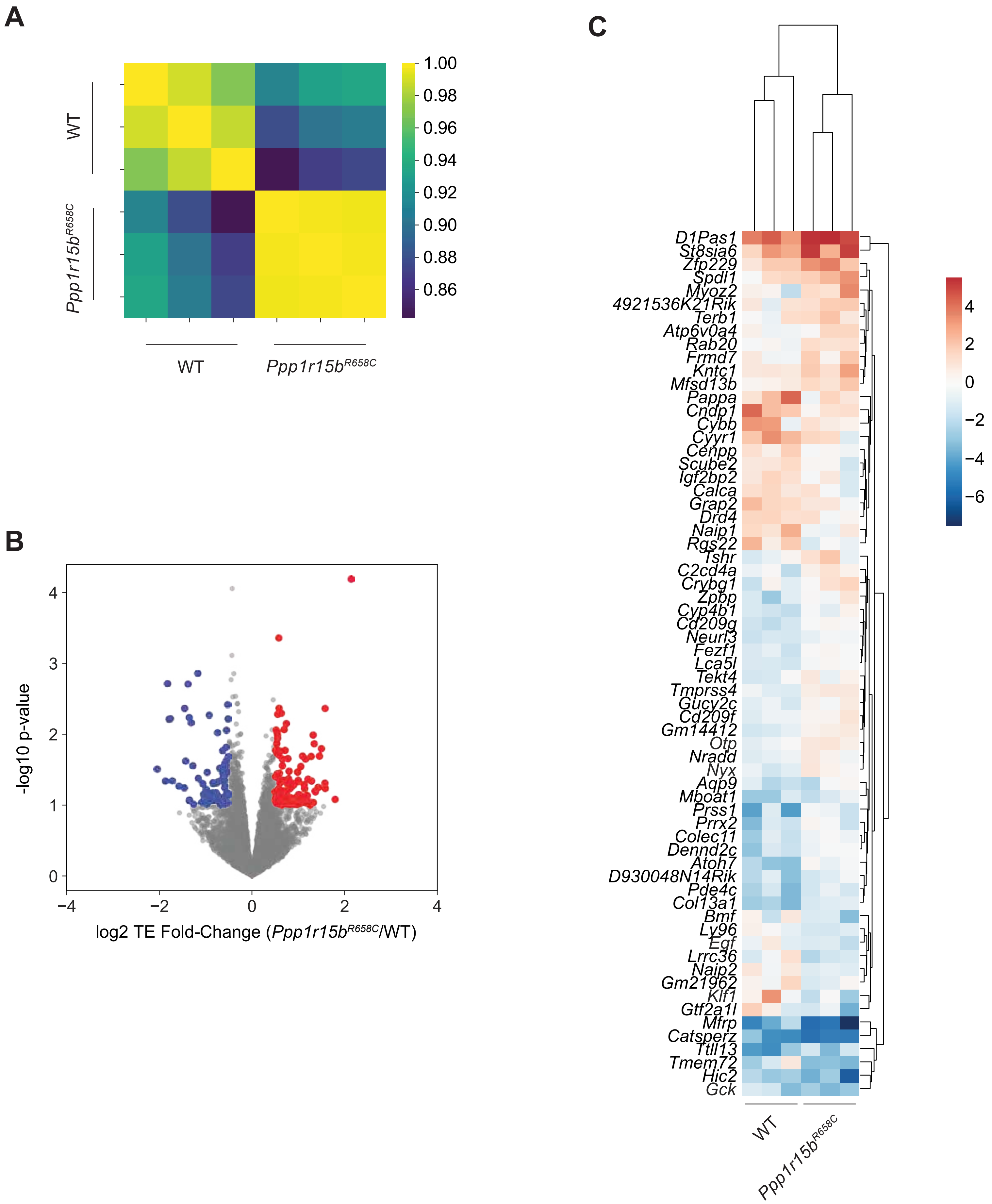


**Fig. S4.** Reineke *et al.*

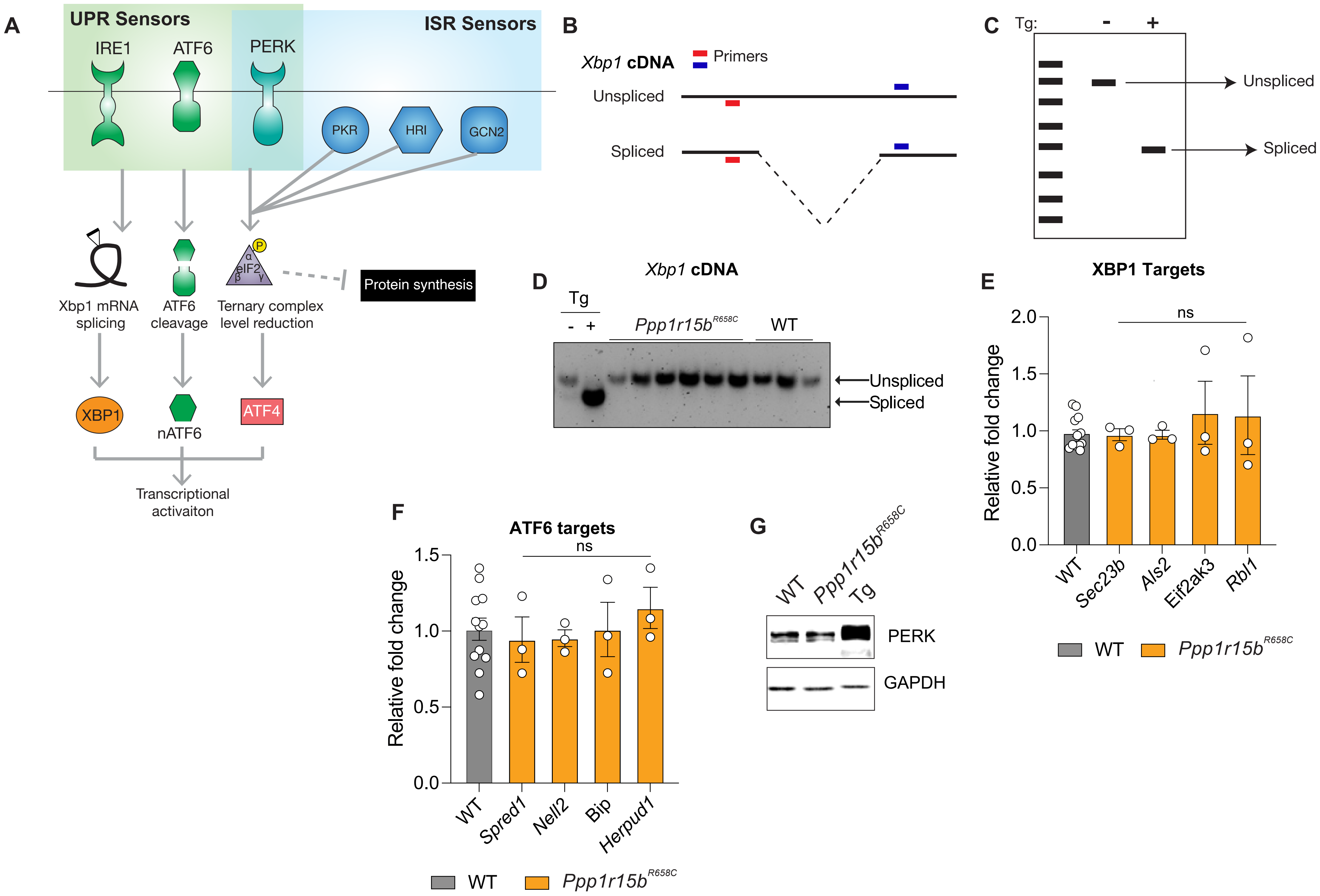


**Fig. S5. Reineke *et al.***

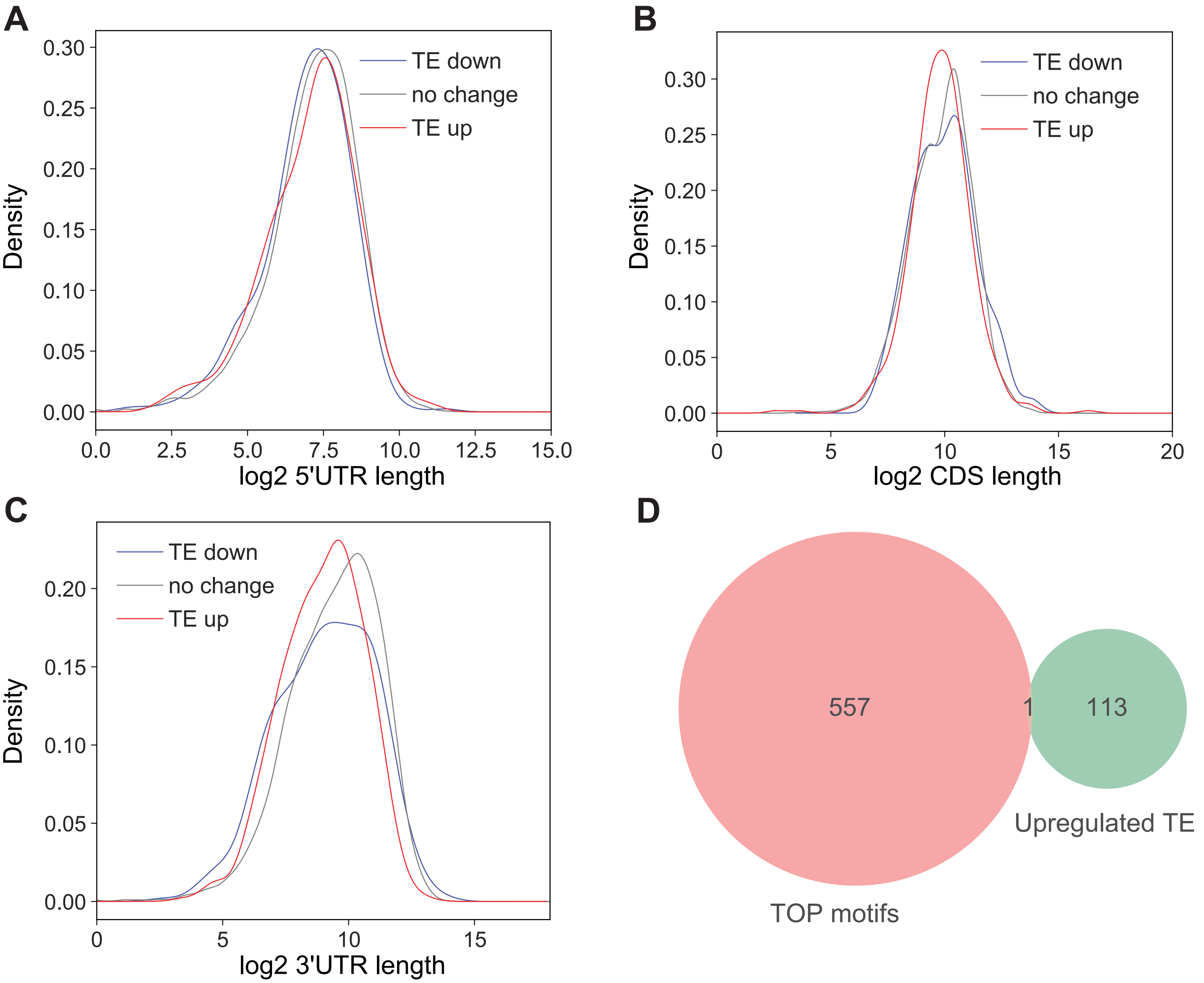


**Fig. S6.** Reineke *et al.*

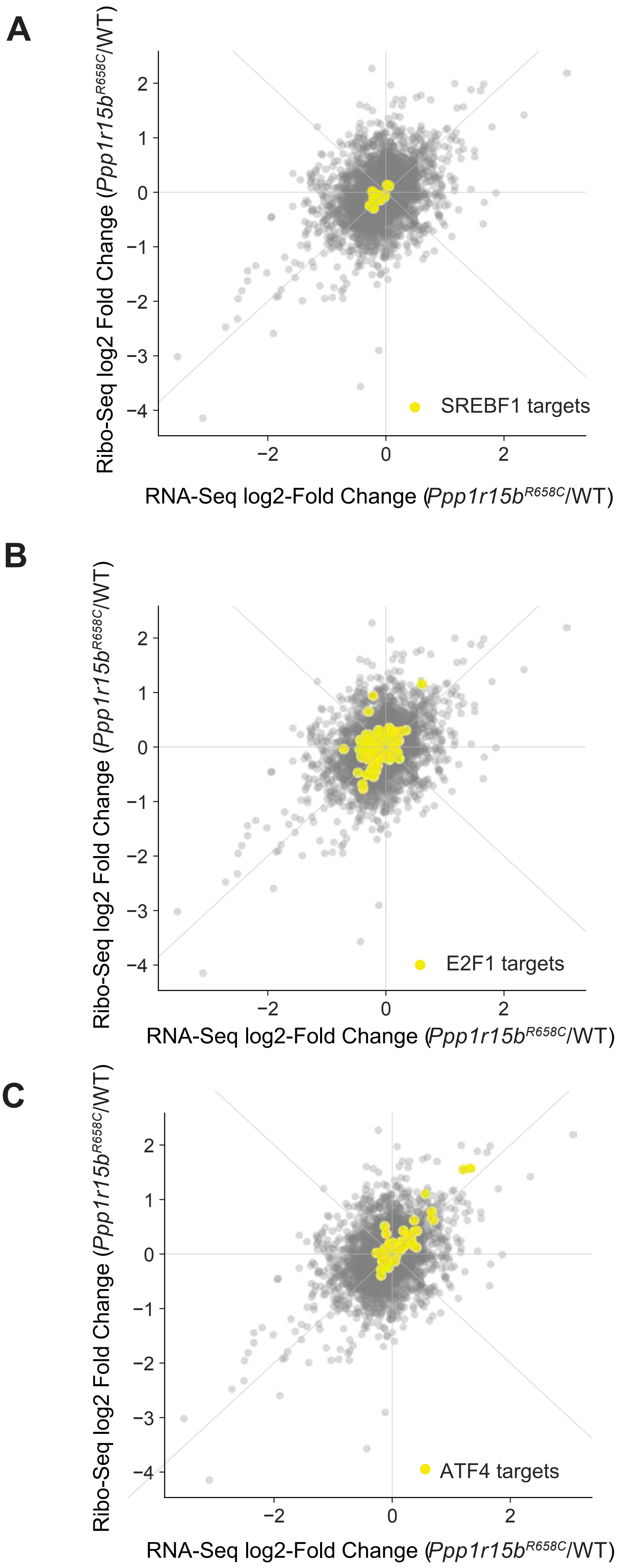


**Fig. S7.** Reineke *et al.*

A

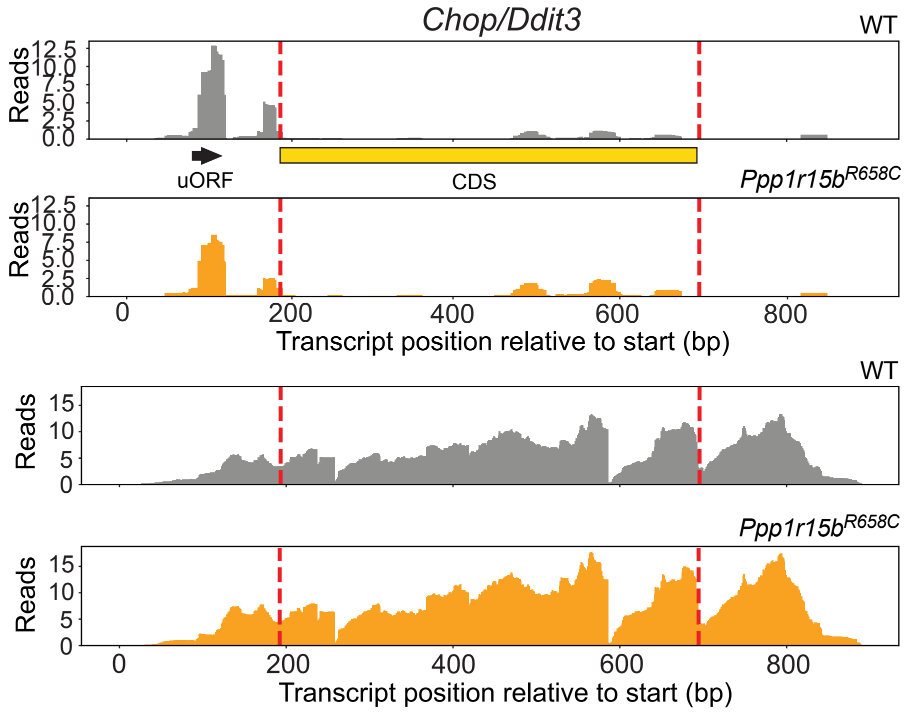


B

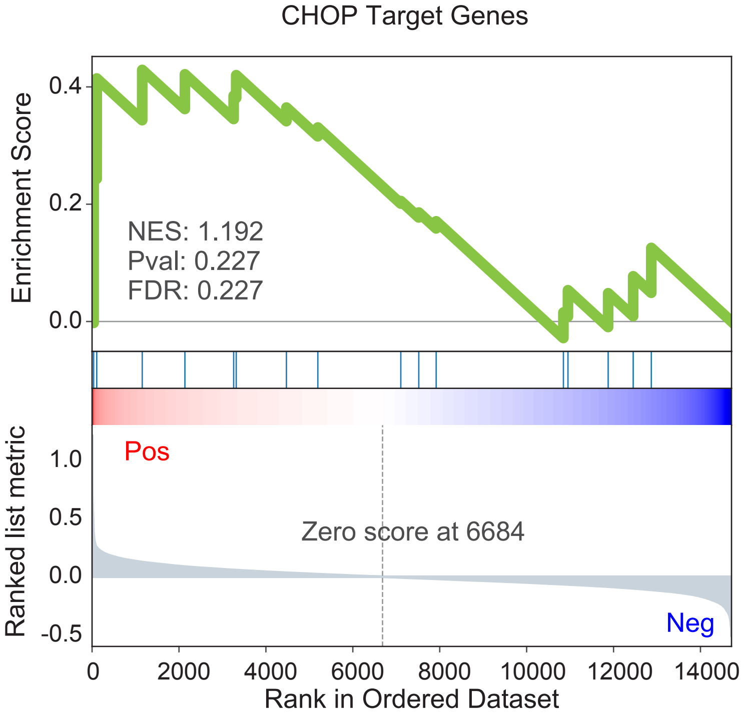


**Fig. S8.** Reineke *et al.*

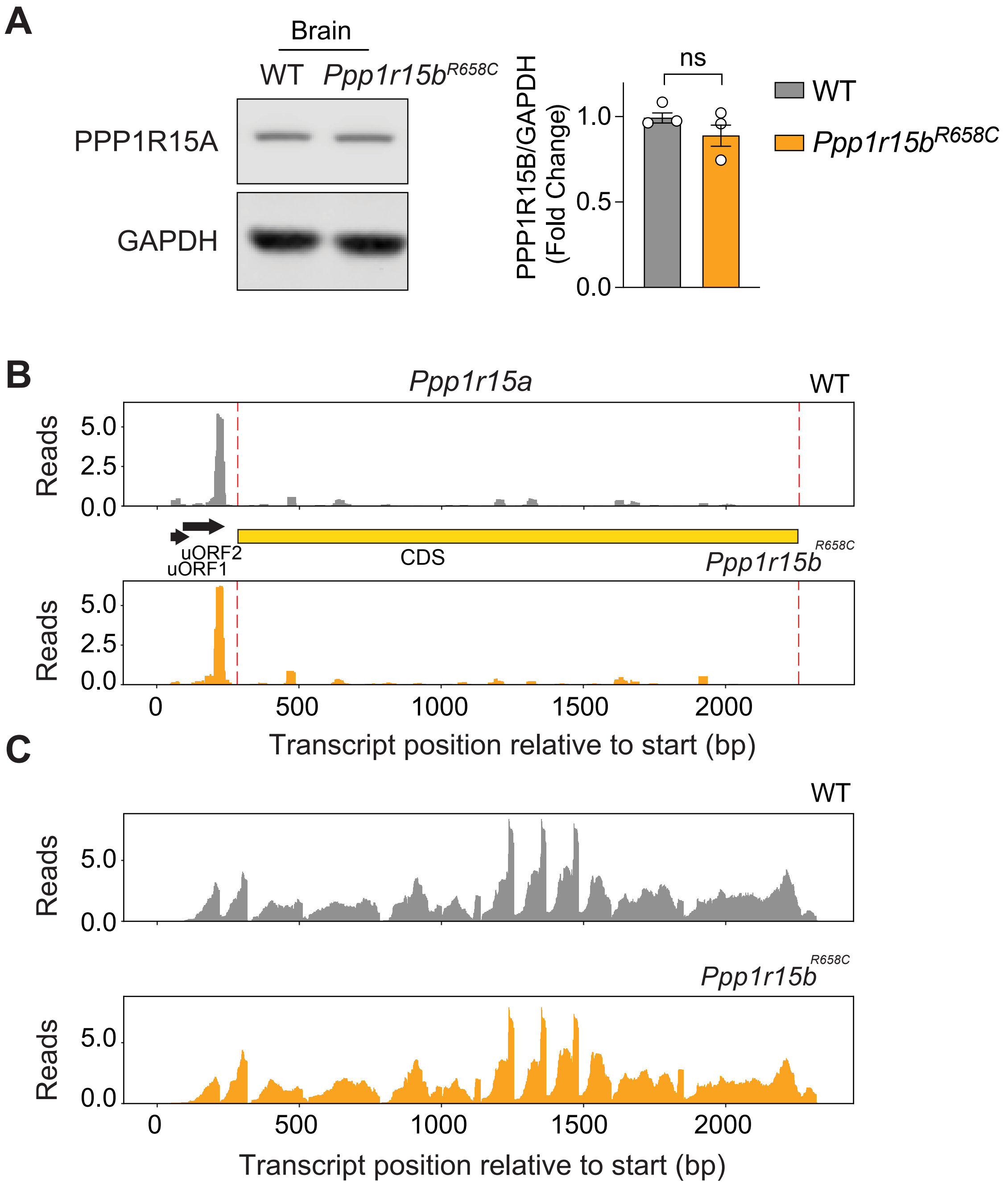


**Fig. S9.** Reineke *et al.*

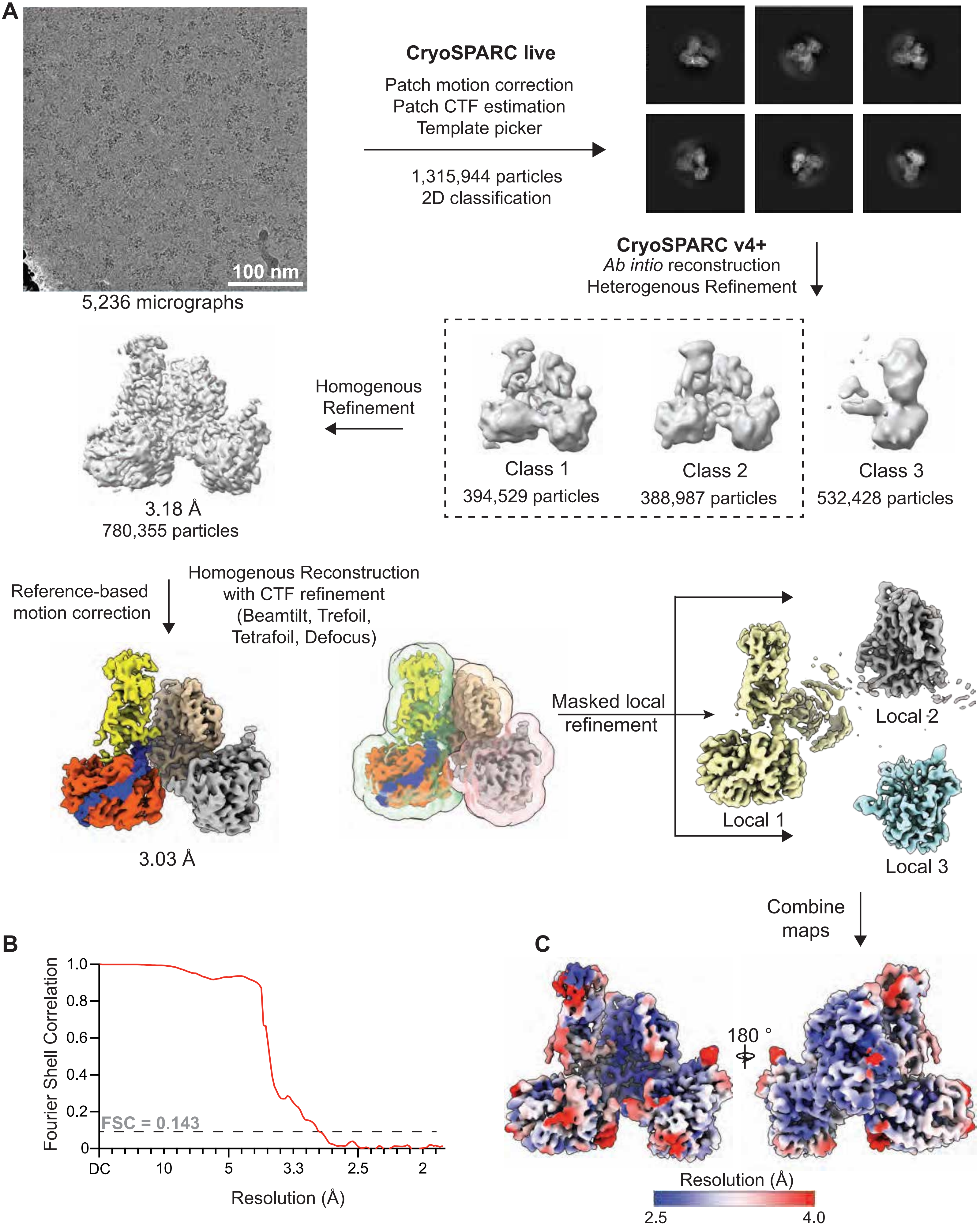


**Fig. S10.** Reineke *et al.*

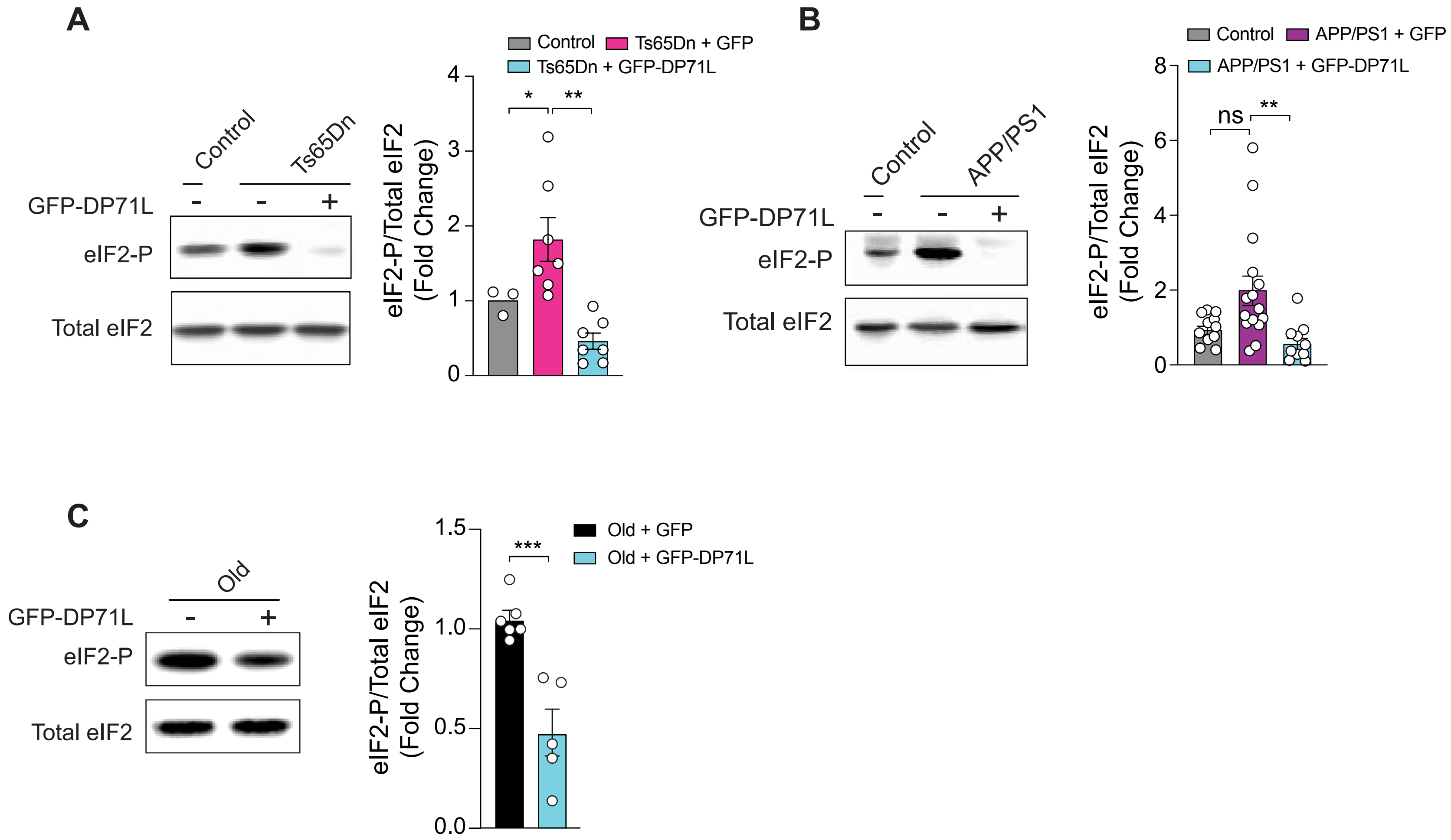


**Fig. S11.** Reineke *et al.*

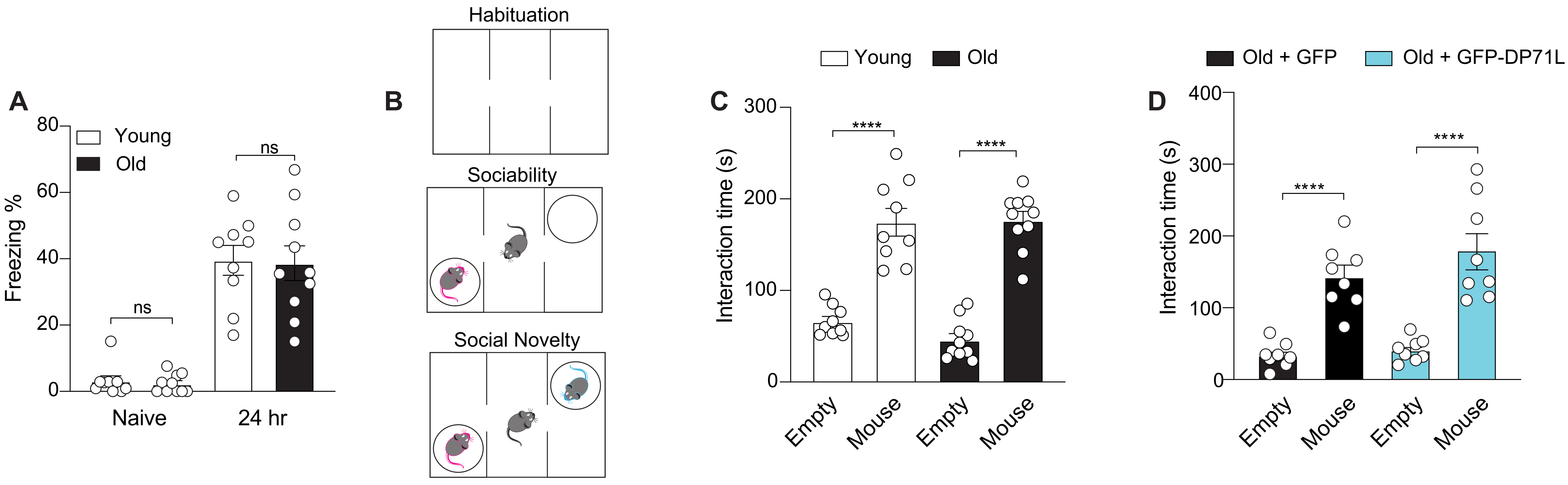


**Fig. S12.** Reineke *et al.*

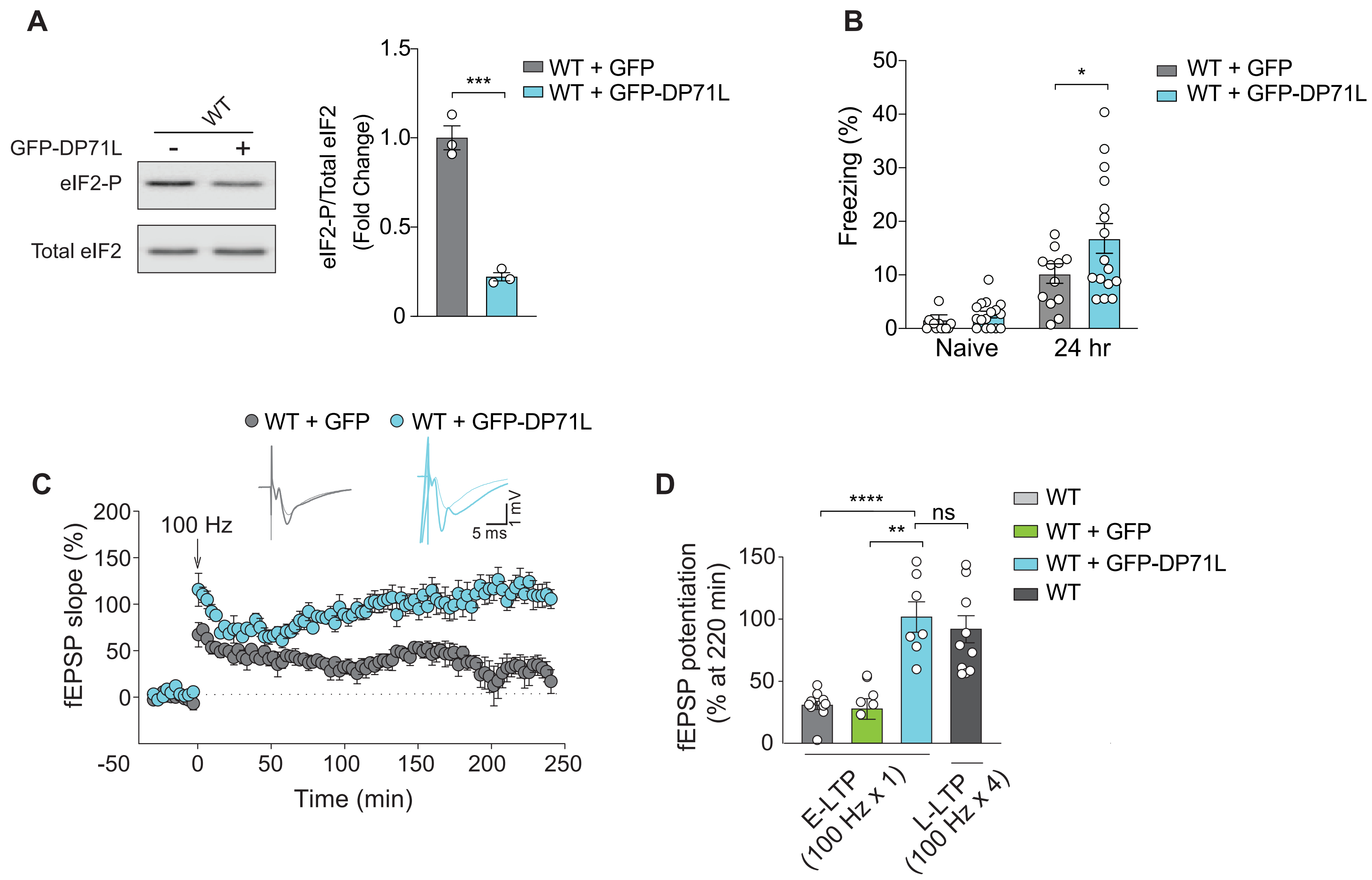


**Fig. S13.** Reineke *et al.*

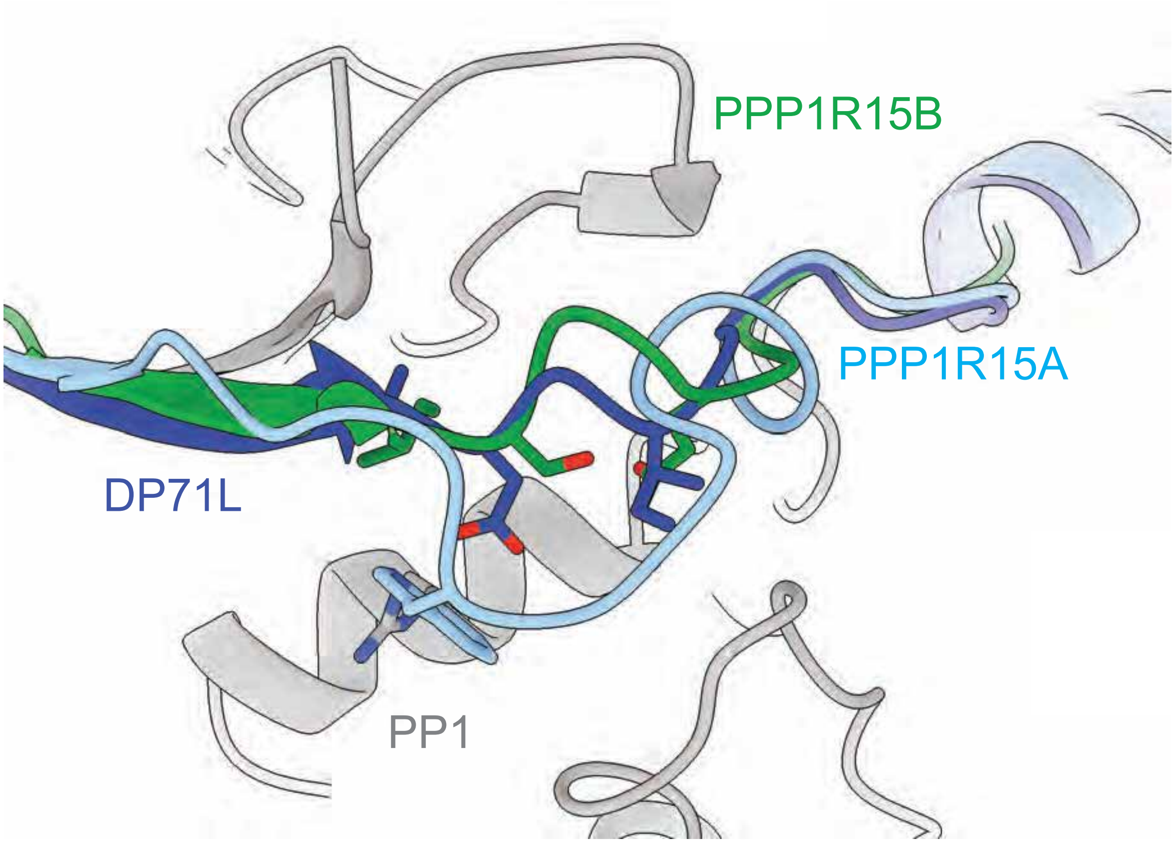


**Fig. S14.** Reineke *et al.*

### Supplementary Figure Legends

**Figure S1. Ternary complex levels are reduced in the brain of *Ppp1r15b*$^{R658C}$ mice.**

(**A**) Schematic representation of the eIF2•eIF2B complex and effects of its inhibition on ternary complex levels.

(**B**) Immunoprecipitation of control (IgG IP) or eIF2 (eIF2β IP) from the brain of WT and *Ppp1r15b*$^{R658C}$ mice followed RT-PCR for the Met-tRNA$_i^{Met}$ and western blotting for eIF2 components.

**Figure S2. Basal synaptic transmission did not differ in WT and *Ppp1r15b*$^{R658C}$ mice.**

(**A**) Paired-pulse facilitation of fEPSPs did not differ between WT ($n$ = 16) and *Ppp1r15b*$^{R658C}$ mice ($n$ = 14), as shown by the plots of the PP ratio (fEPSP2/fEPSP1) for various intervals of paired stimulation.

(**B**) fEPSPs as a function of presynaptic volleys did not differ between WT and *Ppp1r15b*$^{R658C}$ mice ($n$ = 16 for WT, $n$ = 14 for *Ppp1r15b*$^{R658C}$ mice; $R^2$ = 0.47 for WT and 0.52 for *Ppp1r15b*$^{R658C}$ slices).

**Figure S3. Increased inhibitory (but not excitatory) synaptic transmission in CA1 neurons of *Ppp1r15b*$^{R658C}$ mice results from persistent ISR activation.**

(**A-C**) Sample traces (**A**) and summary data show frequency (**B**) and amplitude (**C**) of mEPSCs in CA1 neurons from WT ($n$ = 13) and *Ppp1r15b*$^{R658C}$ mice ($n$ = 15) (frequency: $t$ = 1.06, $P$ = 0.30; amplitude: $t$ = 0.35, $P$ = 0.73).

(**D-F**) Sample traces (**D**) and summary data show frequency (**E**) and amplitude (**F**) of mIPSCs in CA1 neurons from WT ($n$ = 18) and *Ppp1r15b*$^{R658C}$ mice ($n$ = 20) (frequency: $t$ = 5.08, $P < 0.0001$; amplitude: $t$ = 2.14, $P < 0.05$).

(**G-H**) Sample traces (**H**) and summary data show frequency of mIPSCs in CA1 neurons from WT ($n$ = 8), *Ppp1r15b*$^{R658C}$ mice ($n$ = 7) and *Ppp1r15b*$^{R658C}$- *Eif2s1*$^{S/A}$ mice ($n$ = 14) (WT vs. *Ppp1r15b*$^{R658C}$: $t$ =

6.38, $P < 0.0001$; *Ppp1r15b*$^{R658C}$ vs. *Ppp1r15b*$^{R658C}$- *Eif2s1*$^{S/A}$: $t = 5.85$, $P < 0.0001$; WT vs. *Ppp1r15b*$^{R658C}$- *Eif2s1*$^{S/A}$: $t = 0.05$, $P = 0.96$). Data are mean ± SEM. *$P < 0.05$, ****$P < 0.0001$.

**Figure S4. Differentially translated mRNAs in the brain of *Ppp1r15b*$^{R658C}$ mice compared to WT mice.**

(**A**) Heat map of Pearson correlation coefficients of ribosome profiling show high correlation between the three biological replicates from WT and *Ppp1r15b*$^{R658C}$ mice.

(**B**) Volcano plots of up- (red, log2FC ≥ 0.5, p adj ≤ 0.1) and downregulated genes (blue, log2FC ≤ 0.5, p adj ≤ 0.1) identified in the Ribo-seq dataset in the brain of WT and *Ppp1r15b*$^{R658C}$ mice.

(**C**) A heat map showing differentially translated genes in the brain of WT and *Ppp1r15b*$^{R658C}$ mice.

**Figure S5. The UPR is not activated in the brain of *Ppp1r15b*$^{R658C}$ mice.**

(**A**) Schematic of a simplified UPR pathway, and the overlap with the Integrated Stress Response.

(**B-C**) Schematic representation of XBP1 splicing assay. (**B**) RT-PCR was performed to amplify the region of XBP1 mRNA containing the IRE1 splice site. When the UPR is inactive, IRE1 is not activated and XBP1 mRNA is not spliced, resulting in a larger PCR product. When the UPR is active, the XBP1 mRNA is spliced and resulting in a smaller PCR product (**C**).

(**D**) RT-PCR was performed on mRNA isolated from the brains of WT and *Ppp1r15b*$^{R658C}$ mice with primers that specifically recognize spliced vs. unspliced XBP1 mRNA. Fibroblasts were treated with either vehicle (-) or Thapsigargin, Tg (+) as control: treatment with Tg triggers UPR activation and induces XBP1 splicing.

(**E**) RT-PCR was performed on mRNA isolated from the brains of WT ($n = 3$) and *Ppp1r15b*$^{R658C}$ mice ($n = 3$) with primers that amplify XBP1 targets ($F_{4,8} = 0.42$; *Sec23b:* $t = 0.02$, $P > 0.99$; *Als2*: $t = 0.02$, $P > 0.99$; *Eif2ak3*: $t = 0.87$, $P > 0.99$; *Rbl1*: $t = 0.77$, $P > 0.99$).

(**F**) RT-PCR was performed on mRNA brain extracts from WT (*n* = 3) and *Ppp1r15b*$^{R658C}$ mice (*n* = 3) with primers that amplify ATF6 targets ($F_{4,8}$ = 0.27; *Spred1: t* = 0.43, *P* > 0.99; *Nell2*: *t = 0.39, P > 0.99; Bip*: *t = 0.13, P* > 0.99*; Herpud1*: *t = 0.48, P* > 0.99*).* Data are mean ± SEM. ns = not significant.

(**G**) Representative western blots from brain extracts from WT (*n* = 3) and *Ppp1r15b*$^{R658C}$ mice (*n* = 3) using an antibody against PERK. Tg-induced UPR stress activates PERK, as indicated by the upward shift in its migration pattern during gel electrophoresis.

**Figure S6. Features of differentially translated mRNAs in the brains of WT and *Ppp1r15b*$^{R658C}$ mice.**

(**A**) mRNAs with high, low and unchanged TEs (*Ppp1r15b*$^{R658C}$/WT). mRNAs are separated based on the length of their 5' UTR.

(**B**) mRNAs with high, low and unchanged TEs (*Ppp1r15b*$^{R658C}$/WT). mRNAs are separated based on the length of their coding sequence (CDS).

(**C**) mRNAs with high, low and unchanged TEs (*Ppp1r15b*$^{R658C}$/WT). mRNAs are separated based on the length of their 3' UTR.

(**D**) Venn diagram showing overlap between translationally up-regulated mRNAs in *Ppp1r15b*$^{R658C}$ mouse brain and mRNAs containing TOP motifs in their 5'UTR.

**Figure S7. No correlation was found between the genes upregulated in the brains of *Ppp1r15b*$^{R658C}$ mice and those regulated by E2F1 or SREBF1.**

(**A**) SREBF1 target genes are highlighted in differentially expressed genes in Ribo-seq vs. RNA-seq.

(**B**) E2F1 target genes are highlighted in differentially expressed genes in Ribo-seq vs. RNA-seq.

(**C**) ATF4 target genes are highlighted in differentially expressed genes in Ribo-seq vs. RNA-seq.

**Figure S8. Ddit3/CHOP-mediated transcription is not upregulated in the brains of *Ppp1r15b*$^{R658C}$ mice.**

(**A**) Ribosomal read density for Ddit3/CHOP from the ribosome profiling (ribo-seq) dataset (top) and CHOP RNA levels from the RNA sequencing (RNA-seq) dataset (bottom) in WT and *Ppp1r15b*$^{\text{R658C}}$ mice.

(**B**) Gene set enrichment analysis comparing differentially expressed mRNAs in *Ppp1r15b*$^{\text{R658C}}$ mice with the transcriptional targets of CHOP.

**Figure S9. PPP1R15A/GADD34 is not up regulated in the brains of *Ppp1r15b*$^{\text{R658C}}$ mice.**

(**A**) Representative Western blot (left) and quantification (right) for PPP1R15A/GADD34 protein in the brain of WT and *Ppp1r15b*$^{R658C}$ mice ($n$ = 4 per group, $t$ = 1.56, $P$ = 0.17)

(**B**) Ribo-seq reads on the *Ppp1R15a*/*Gadd34* mRNA in WT (top) and *Ppp1r15b*$^{R658C}$ (bottom) animals.

(**C**) Reads from RNA-seq mapped along the *Ppp1R15a*/*Gadd34* mRNA in WT (top) and *Ppp1r15b*$^{R658C}$ (bottom) animals. Data are mean ± SEM. ns = not significant

**Figure S10. Cryo-EM data acquisition and analysis workflow.**

(**A**) An overview of the data processing steps used to generate the final 3D reconstruction of DP71L in complex with p-eIF2$\alpha$NTD, PP1A, G-actin, and DNAseI.

(**B**) Fourier shell correlation (FSC) curves of the consensus and locally refined maps generated using the cryoSPARC auto-generated tight mask followed by high resolution noise substitution.

(**C**) The combined map of the three locally refined structures colored by local resolution measured using an FSC threshold of 0.143.

**Figure S11. DP71L expression suppresses the ISR in the brains of various models of cognitive dysfunction.**

(**A**) Representative Western blot (left) and quantification (right) of eIF2-P levels in hippocampal extracts from control mice (n = 3) and Ts65Dn mice injected with either AAV-GFP (*n* = 7) or AAV-GFP-DP71L (*n* = 7) [$F_{2,8}$ = 16.33; Control vs. Ts65Dn+GFP: *t* = 3.20, *P* < 0.05; Ts65Dn+GFP vs. Ts65Dn+DP71L: *t* = 5.56, *P* < 0.01; Control vs. Ts65Dn+DP71L: *t* = 0.72, *P* > 0.99].

(**B**) Representative Western blot (left) and quantification (right) of eIF2-P levels in hippocampal extracts from control mice (*n* = 11) or APP/PS1 mice injected with either AAV-GFP (*n* = 15) or AAV-GFP-DP71L (*n* = 10) [$F_{2,10}$ = 7.47; Control vs. APP/PS1 + GFP: *t* = 1.69, *P* = 0.36; APP/PS1 + GFP vs. APP/PS1 + DP71L: *t* = 3.83, *P* < 0.01; Control vs. APP/PS1 + DP71L: *t* = 0.73, *P* > 0.99].

(**C**) Representative Western blot (left) and quantification (right) of eIF2-P levels in hippocampal extracts from old mice injected with either AAV-GFP (*n* = 6) or AAV-GFP-DP71L (*n* = 5) [Old + GFP vs. Old + GFP-DP71L: *t* = 4.89, *P* < 0.001]. Data are mean ± SEM. **P* < 0.05, ***P* < 0.01, ****P* < 0.001, ns = not significant.

**Figure S12. Long-term fear memory and sociability is similar in young and old mice.**

(**A**) Long-term contextual fear memory in control in young (*n* = 9) and old animals (*n* = 10) [$F_{3,25}$ = 25.41; Young vs. Old: *t* = 0.10, *P* > 0.99].

(**B**) Schematic of sociability and social novelty behavioral paradigms.

(**C**) Three chamber sociability in young (*n* = 9) and old (*n* = 10) mice [$F_{3,25}$ = 188.40; Young Empty vs. Young Mouse: *t* = 14.23, *P* < 0.0001; Old Empty vs. Old Mouse: *t* = 18.88, *P* < 0.0001].

(**D**) Sociability in old mice injected with either AAV-GFP (n = 8) or AAV-GFP-DP71L (n = 8) [$F_{3,21}$ = 47.06; Old + GFP Empty vs. Old + GFP Mouse: *t* = 7.33, *P* < 0.0001; Old + DP71L Empty vs. Old + DP71L Mouse: *t* = 9.08, *P* < 0.0001]. Data are mean ± SEM. *****P* < 0.0001, ns = not significant.

**Figure S13. DP71L facilitates long term memory formation and long-lasting changes in synaptic function in normal healthy mice.**

(**A**) Representative Western blot (left) and quantification (right) of eIF2-P levels in the brain of WT mice injected with either AAV-GFP ($n$ = 3) or AAV-GFP-DP71L ($n$ = 3) [WT + GFP vs. WT + DP71L: $t$ = 9.80, $P$ < 0.001].

(**B**) Long-term fear memory in WT mice injected with either AAV-GFP ($n$ = 12) or AAV-GFP-DP71L ($n$ = 16) [$F_{1,54}$ = 2.36, $t$ = 2.53, $P$ < 0.05]. Freezing times were recorded before conditioning (naive, during 2 min period) and then 24 hr after training (during 5 min period). Mice were subjected to a weak fear conditioning protocol (a single pairing of a tone with a 0.35 mA foot shock).

(**C**) A single high-frequency train (100 Hz for 1 s) elicited a short-lasting early LTP (E-LTP) in in WT mice injected with AAV-GFP ($n$ = 6) but generated a sustained late LTP (L-LTP) in WT mice injected with AAV-GFP-DP71L ($n$ = 6) [at 220 min: $H$ = 9.60, $Q$ = 3.098, $P$ < 0.001].

(**D**) Comparison of the magnitude of the synaptic potentiation elicited with an L-LTP-inducing protocol (four trains at 100Hz, separated by 5 min) in slices from WT mice ($n$ = 11) vs. an E-LTP-inducing protocol (a single train at 100 Hz) induced in slices from WT mice injected with either AAV-GFP ($n$ = 6) or AAV-GFP-DP71L ($n$ = 7) [$F_{3,20}$ = 21.39; WT vs. WT + GFP-DP71L: $t$ = 6.47, $P$ < 0.0001; WT + GFP vs. WT + GFP-DP71L: $t$ = 4.30, $P$ < 0.01; WT + GFP-DP71L 100 Hz x 1 vs. WT 100 Hz x 4: $t$ = 0.87, $P$ > 0.99]. Data are mean ± SEM. *$P$ < 0.05, **$P$ < 0.01, ***$P$ < 0.001, ****$P$ < 0.0001, ns = not significant.

**Figure S14. Structural modeling of the linker region of the mammalian (PPP1R15A and PPP1R15B) and viral (DP71L) phosphatase cofactors.**

In contrast to DP71L, structural modeling of the linker region in PPP1R15A and PPP1R15B reveals an enlarged bulge and a loss of hydrogen bonding, which results in a reduced binding affinity for PP1.
